# Aberrant regulation of serine metabolism drives extracellular vesicle release and cancer progression

**DOI:** 10.1101/2022.05.10.491299

**Authors:** Tomofumi Yamamoto, Jun Nakayama, Fumihiko Urabe, Kagenori Ito, Nao Nishida-Aoki, Masami Kitagawa, Akira Yokoi, Masahiko Kuroda, Yutaka Hattori, Yusuke Yamamoto, Takahiro Ochiya

**Author notes:** corresponding authors **Corresponding to:** Takahiro Ochiya, Ph.D. Professor, Department of Molecular and Cellular Medicine, Institute of Medical Science, Tokyo Medical University 6-7-1, Nishishinjyuku, Shinjyuku-ku, Tokyo 160-0023, Japan E-mail, Yusuke Yamamoto, Ph.D. Unit leader, Laboratory of Integrative Oncology, National Cancer Center Research Institute, Tokyo 5-1-1, Tsukiji, Chuo-ku, Tokyo 104-0045, Japan. equally contributed authors (Tomofumi Yamamoto, Jun Nakayama).

## Abstract

Cancer cells secrete extracellular vesicles (EVs) to regulate cells in the tumor microenvironment to benefit their own growth and survive in the patient’s body. Although emerging evidence has demonstrated the molecular mechanisms of EV release, regulating cancer-specific EV secretion remains challenging. In this study, we applied a microRNA library to reveal the universal mechanisms of EV secretion from cancer cells. Here, we identified miR-891b and its direct target gene, phosphoserine aminotransferase 1 (PSAT1), which promotes EV secretion through the serine-ceramide synthesis pathway. Inhibition of PSAT1 affected EV secretion in multiple types of cancer, suggesting that the miR-891b/PSAT1 axis shares a common mechanism of EV secretion from cancer cells. Interestingly, aberrant PSAT1 expression also regulated cancer metastasis via EV secretion. Our data link between the PSAT1-controlled EV secretion mechanism and cancer metastasis and show the potential of this mechanism as a therapeutic target in multiple types of cancer.

## Introduction

Exosomes are small EVs (small extracellular vesicles: sEVs) with a diameter of approximately 50–150 nm (1–2). In general, cancer cells secrete a large number of exosomes comparing to normal cells (3) to modify the tumor microenvironment and support the growth of tumor cells (4–6). Therefore, inhibiting exosome secretion by cancer cells will have therapeutic benefits to be a less tumor-promoting environment (7).

EV secretion can be generally divided into two pathways: the endosomal sorting complex required for transport complex (ESCRT)-dependent pathway and the ESCRT-independent pathway. Exosomes are formed by inward budding into early endosomes, which becomes multivesicular bodies (MVBs), and released into the extracellular environment through MVB fusion to the cellular membrane (8–10). ALIX and TSG101 are proteins related to the ESCRT-dependent endosomal pathway, while the ESCRT-independent pathway includes nSMase2, which is involved in ceramide synthesis (3, 11–12). One of the reasons for increase of secretion is that oncogenic signaling, such as signaling through MYC and RAS, promotes EV release (13). Recently, we reported that miR-26a regulates EV secretion from PC-3M prostate cancer cells by targeting SHC4, PFDN4 and CHORDC1, resulting in tumor suppression (14). Furthermore, EV secretion was regulated via miR-1908/spermidine synthase (SRM) in 22Rv1 prostate cancer cells (15). Thus, cancer cells seem to utilize a different EV secretion machinery from non-cancerous cells. Several small molecules were reported to regulate EV secretion. For example, P2RX7 inhibitor, which suppresses the disease-related EV secretion, leading to attenuating the disease phenotype (16). GW4869 inhibits cancer cell-derived EV secretion; however, it also affected EV secretion from normal cells (17). Additionally, the cargo of EVs is important for cancer progression (18–20).

Our previous study reported that we bypassed the complicated EV purification process and examined the effect of individual miRNA on EV secretion in a high-throughput manner by an ExoScreen method (21). To further elucidate the universal molecular mechanisms of EV secretion, we performed a microRNA (miRNA) library screen of cell lines from colorectal and lung cancer. In the process of performing the miRNA library screen, we used the ExoScreen assay to systematically quantify the EV amount. From the miRNA-based screen, we identified miR-891b, which suppressed the EV secretion in cancer cell lines. Additionally, as a direct target gene of miR-891b, we identified phosphoserine-aminotransferase 1 (PSAT1), which positively regulated EV secretion via the serine de novo synthesis pathway. Our findings revealed the molecular mechanism of PSAT1-controlled EV secretion in multiple types of cancer cells and demonstrated the contribution of PSAT1 to cancer metastasis in breast cancer models.

## Results

### miR-891b suppresses EV secretion from colon and lung cancer cell lines

To identify novel molecular mechanisms of EV secretion, we performed a miRNA-based screen of the HCT116 colon cancer and A549 lung adenocarcinoma cell lines using a miRNA mimic library (1728 miRNAs in total). The amount of EVs in the conditioned medium collected from miRNA-transfected cells was quantified by the ExoScreen assay, which we previously developed as an ultrasensitive EV detection method (21). This method enables us to detect EVs without any purification steps. Two types of antibodies are used to capture EVs and then, are detected by photosensitizer-beads. To exclude miRNAs that decrease the EV number through cell death, we assessed the cellular proliferation simultaneously **(Figure 1A)**. To confirm EV secretion, the EV size and particle number were evaluated by nanoparticle tracking analysis (NTA), and surface markers of EVs, i.e., CD9 and CD63, were verified by Western blot analysis **(Figures 1B and 1C)**. Among the genes regulating EV secretion, TSG101 was used as a positive control for EV secretion screening **(Figure S1A)**. From the library of 1728 miRNAs, we excluded miRNAs that affected cell proliferation (CCK-8 signal normalized to that in negative control cells= < 0.8 or > 1.2) **(Figure 1D)**. After selection, the inhibitory effects of the miRNAs (based on the z-scores) on EV secretion were quantified in both cell lines **(Figure 1E)**. As shown in **Figure 1F**, the top 10 miRNAs with the highest z-scores were selected. Two miRNAs, miR-891b and miR-526a, were selected in both cell lines (marked in red). We validated the efficacy of the selected miRNAs **(Figure S1B)**, and focused on miR-891b for further analysis as the most effective miRNA. miR-891 decreased the secretion of both CD9-positive EVs and CD63-positive EVs to 40-60% based on the ExoScreen assay of conditioned medium and isolated EVs by ultracentrifugation **(Figure S1C)**. The decrease of EV secretion was further verified by NTA **(Figures S1D and S1E)**. Additionally, high expression of miR-891b was significantly correlated with better prognosis in lung adenocarcinoma, although the correlation was not statistically significant in colorectal cancer **(Figure S1F)**.

**Fig. 1.**
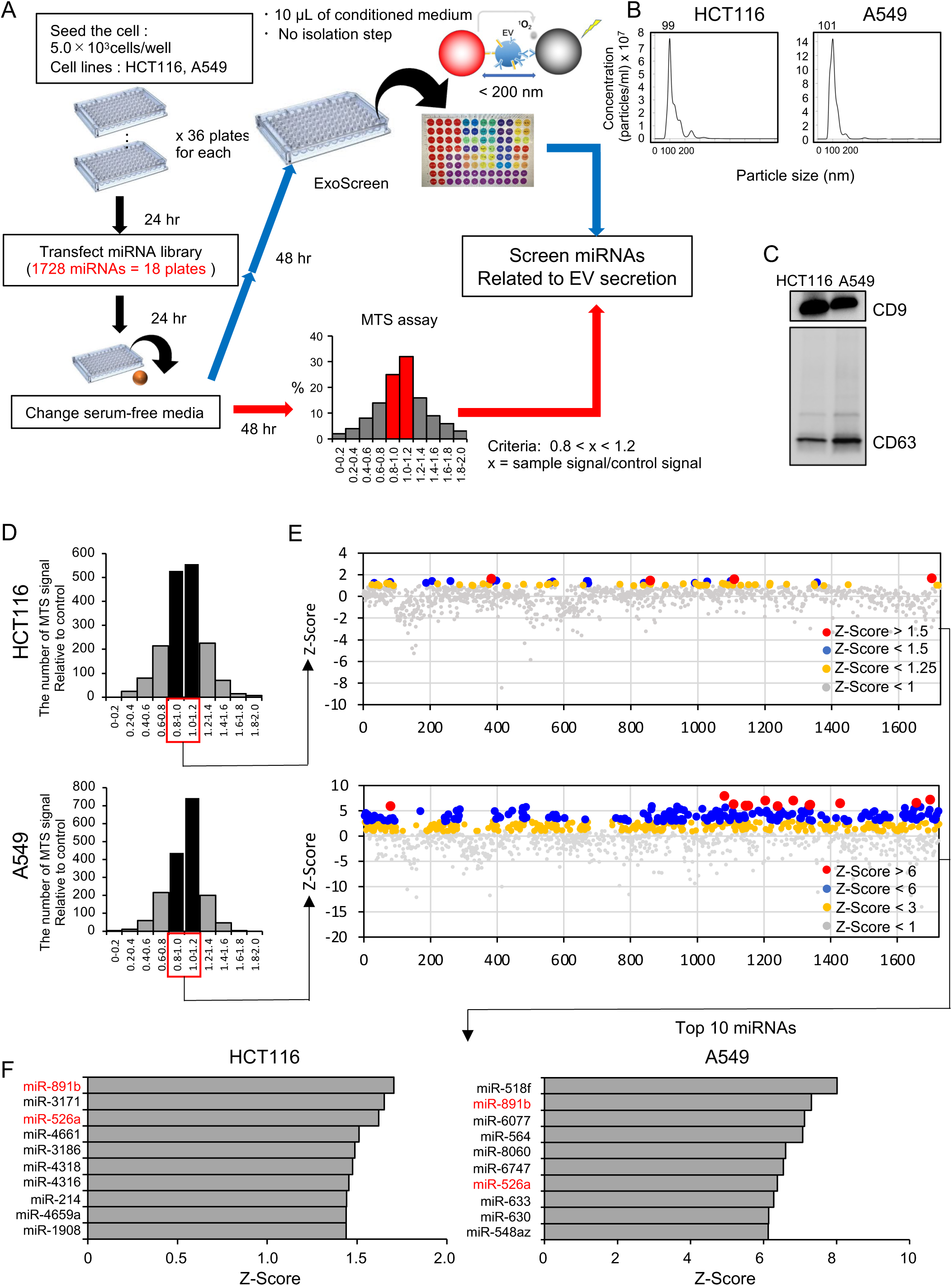
miRNA screening identified miR-891b as being responsible for EV secretion. **A.** A schematic of the miRNA-based screening approach for identifying EV secretion-related miRNAs. **B.** NTA of EVs derived from HCT116 and A549 cells. **C.** Western blot analysis of the EV surface markers CD9 and CD63 in EV samples collected from HCT116 and A549 cells. EVs were loaded at 500 ng protein/lane. **D.** Relative cell viability after introducing miRNA mimics was evaluated with the MTS assay. The signals were normalized to those in the negative control group. miRNAs with a signal intensity of < 0.8 or > 1.2 were excluded. **E.** Scatter plots of EV secretion levels of cancer cells transfected with each miRNA. Top: HCT116; bottom: A549. Relative EV secretion changes were calculated as Z-scores. Only miRNAs with a low impact on cellular viability are shown (0.8 < viability rate< 1.2). **F.** Top 10 miRNAs with high Z-scores for inhibition of EV secretion.

### PSAT1 is one of the direct target genes of miR-891b

To investigate the molecular mechanism of miR-891b in EV secretion, we identified genes that was suppressed by miR-891b and related to EV secretion. We performed transcriptome analysis using HCT116 and A549 cells transfected with either a miR-891b mimic or a control mimic. In the results indicated that 2,221 and 4,711 genes were downregulated (fold change < -1.5) in miR-891b mimic-transfected HCT116 and A549 cells, respectively. We also prepared a gene list that was predicted to be miR-891b target genes according to the TargetScan database (2,618 genes). Comparing the three gene lists, we selected 82 genes overlapped among them (**Figure 2A and Table S1)**. Based on a literature search, we selected 45 of these 82 genes that are likely related to endosome organization and lipogenesis. Then, we examined whether silencing of these 45 genes suppressed EV secretion, consistent with the effect of miR-891b transfection (**Figure 2B**). The ExoScreen assay showed that attenuating PSAT1 gene expression significantly decreased CD9-positive and CD63-positive EVs in both cell lines (**Figures 2C and 2D**). The number of EV particles secreted from PSAT1-knockdown cells was also decreased significantly (**Figures 2E and 2F**). Because a mixture of 4 different siRNA sequences targeting PSAT1 for knockdown was used for this screen, we also examined the effects of the individual PSAT1 siRNAs on EV secretion. Although the effect of siPSAT1-4 was not statistically significant, transfection of the other 3 siRNAs resulted in a significant decrease in EV secretion **(Figures S2A and S2B)**. In addition to PSAT1, we identified other candidate genes, PDXK, RAB31, and VPS37B, that decreased EV secretion by the ExoScreen **(Figure S2C)**. Although the effect of VPS37B on EV secretion was not verified by NTA, silencing of PDXK and RAB31 decreased the particle number of EVs **(Figure S2D)**.

**Fig. 2.**
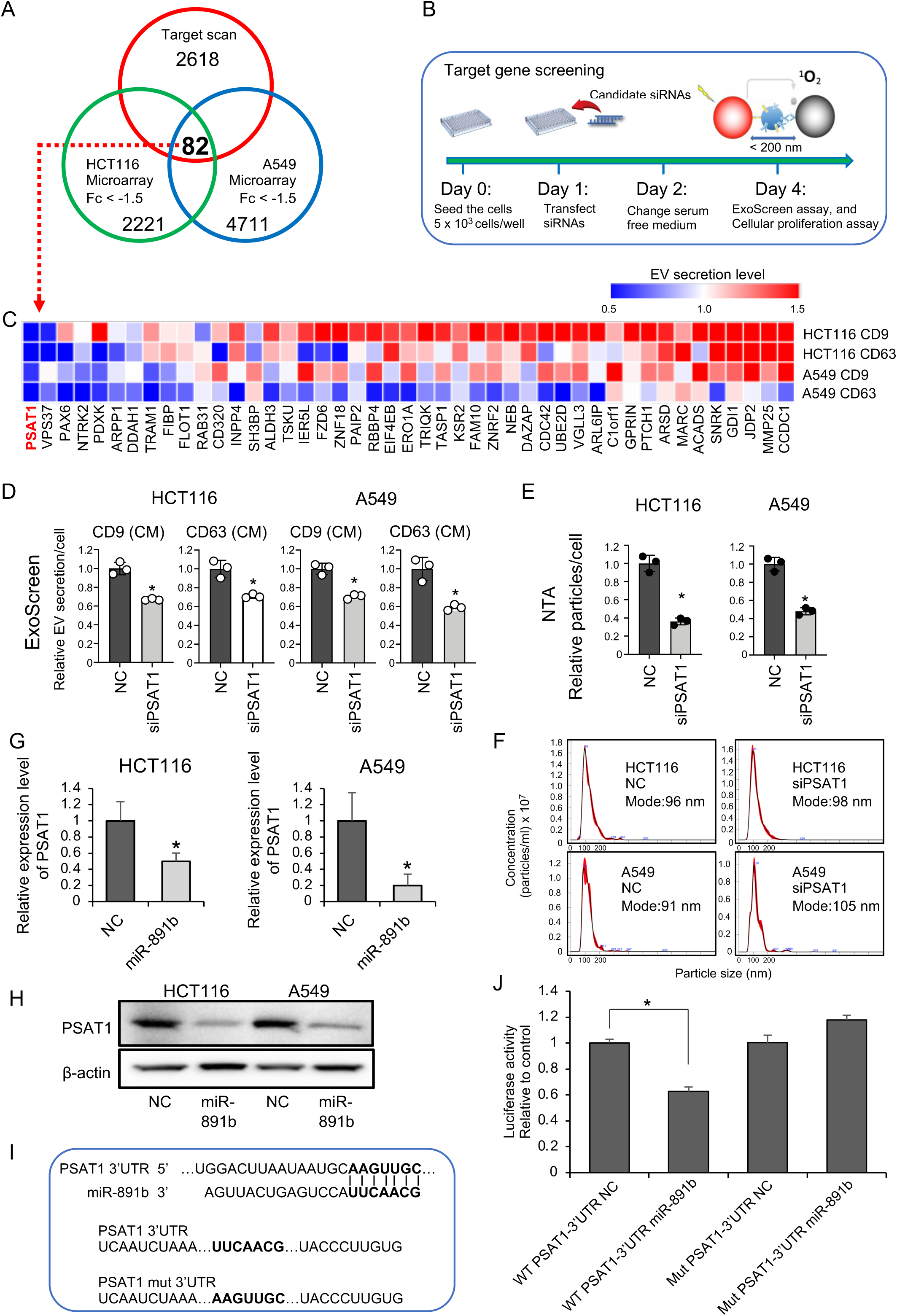
Screening of miR-891b target genes. **A.** Venn diagram showing the overlaps between genes downregulated by miR-891b transfection in HCT116 and A549 cells (Fc < -1.5) and target genes of miR-891b predicted by TargetScan. **B.** Schematic of the approach for miR-891b target gene screening. After siRNAs were transfected into HCT116 or A549 cells, the amount of secreted EVs (both CD9- and CD63-positive) was measured by an ExoScreen assay. **C.** Heatmap showing EV secretion levels of cells with gene silencing. **D.** EV secretion levels after siPSAT1 transfection. EV secretion was measured by the ExoScreen assay based on the surface CD9 or CD63 levels. The values are presented as the means ± SDs (n=3). *p<0.05 by the Student’s t test. **E.** NTA of EVs collected from siPSAT1-transfected cancer cells. The particle count was normalized by the cell number in the collected conditioned medium. The values are presented as the means ± SDs. *p< 0.05 by the Student’s t test. **F**. Representative images of NTA of EVs collected from siPSAT1-transfected cancer cells. **G.** PSAT1 expression levels in miR-891b-transfected cancer cells. The relative expression levels of PSAT1 were normalized to β-actin expression. The values are presented as the means ± SDs (n=3). *p<0.05 by Student t test. **H.** Western blot analysis of PSAT1 after miR-891b was transfected. Each lane was loaded with 15 μg of protein. **I.** Summary of miR-891b target sites and mutated sites (shown in bold) in the 3’UTRs of PSAT1. **J.** The luciferase reporter assay with the wild-type and mutant PSAT1 3’UTR vectors cotransfected with miR-891b. The values are presented as the means ± SDs (n=4, *p<0.05, Student’s t test)

Next, we examined whether PSAT1 is a direct target of miR-891b. The PSAT1 mRNA expression and protein amount were significantly decreased after transfection with the miR-891b mimic, as confirmed by qRT–PCR (**Figure 2G**) and Western blotting (**Figure 2H**). Then, we conducted a luciferase 3’UTR reporter assay to examine whether miR-891b can directly bind to the 3’UTR of PSAT1. We constructed assay vectors with several sequence variations, including the predicted miR-891b target sequences (**Figure 2I**). The assay revealed that wild-type PSAT1 3’UTR, but not other variants, was suppressed by miR-891b (**Figure 2J**). These results revealed that PSAT1 is directly targeted by miR-891b.

To further confirm the effect of PSAT1 on EV secretion, we performed a rescue experiment. We established PSAT1-knockout (KO) A549 cells by CRISPR-Cas9 and then reintroduced PSAT1 into the cells **(Figures S2E and S2F)**. Decreased EV secretion was observed in PSAT1 KO cells, and when PSAT1 overexpressed in the PSAT1 KO cells, the amount of EV secretion was significantly restored. Because PSAT1 is related to a serine synthesis pathway, cellular serine levels were measured in this experiment. Notably, EV secretion and serine levels were highly correlated **(Figure S2G)**. Additionally, we investigated the effect of coexpression of miR-891b and PSAT1 on EV secretion. When PSAT1 was overexpressed in miR-891b-transfected cells, the EV particle number showed an increasing trend in both the HCT116 and A549 cell lines **(Figures S2H and S2I)**. We also examined the effect of PSAT1 knockdown on microvesicle secretion. The microvesicles were isolated from conditioned medium from HCT116 and A549 cells by centrifugation (10,000 g for 40 min). Large size of particles was observed by NTA (**Figure S3A**). The particle numbers were also measured by NTA, and the results showed no significant difference between the PSAT1 knockdown and control samples, indicating that PSAT1 is not related to the secretion of microvesicles (**Figure S3B**).

### The serine synthesis-ceramide pathway contributes to EV secretion in cancer via PSAT1

Since PSAT1 is a famous regulator gene in the serine synthesis pathway **(Figure 3A)** and PSAT1 levels were correlated with cellular serine levels (**Figure S2F and S2G**), we investigated the importance of serine synthesis in EV biogenesis. A serine-free medium (Minimum Essential Media, MEM) was used for examination of the impact of serine on EV secretion **(Figure 3B)**. As expected, while siPSAT1-transfected cells grew at a similar speed in a normal culture medium, siPSAT1-transfected cancer cells grew slower in the serine-depleted medium (**Figure S4A**). In the absence of both serine and FBS in the culture medium for collecting EVs, we observed large-size EVs, which were considered as apoptotic bodies from dying cells (**Figure S4B**). Although the dying cells and secreted apoptotic bodies might affect the EV particles, the number of secreted EV particles significantly increased in several cancer cell lines by the addition of serine (**Figure S4C**). We next investigated whether serine supplementation can rescue EV secretion in siPSAT1-transfected cancer cells. As shown in the EV quantification results obtained by ExoScreen **(Figures 3C and 3D)** and NTA **(Figures 3E-3H)**, the serine-free condition decreased EV secretion level to 30-40% in PSAT1 knockdown cells when normalized by cell number, and supplementing serine (4 mM) restored EV secretion to a similar level to control samples. Additionally, because serine is one of the components of ceramide (0.5 μM), we supplemented ceramide in PSAT1 knockdown cells and examined EV secretion. Similar to serine supplementation, the ceramide supplementation restored EV secretion in PSAT1-knockdown cells (**Figure S4D**). These data indicated that PSAT1-mediated serine-ceramide synthesis is responsible for EV secretion.

**Fig. 3.**
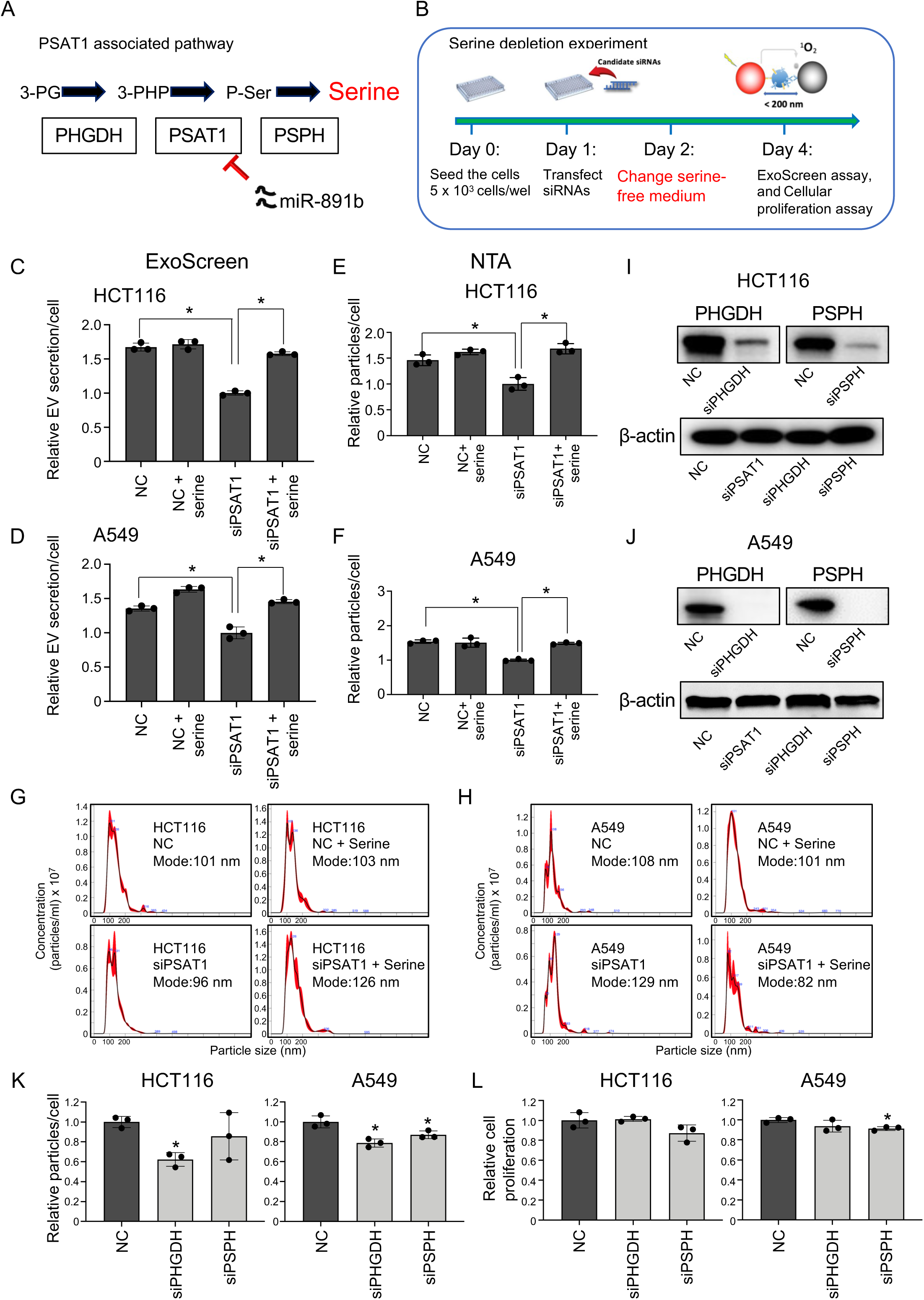
The de novo serine synthesis pathway affects EV secretion in cancer cell lines. **A.** Schematic of the de novo serine synthesis pathway. **B.** Schematic for the serine supplementation experiments with EVs. **C, D.** Rescue experiments were performed to determine the effect of siPSAT1 upon supplementation with serine. After transfection with siPSAT1, serine (4 mM) was added to the serine-free medium. EV secretion was measured by the ExoScreen method. The values are presented as the means ± SDs (n=3). *p<0.05 by one-way ANOVA with post-hoc Tukey’s post hoc test. **E, F.** NTA of EVs secreted in the rescue experiments with serine supplementation. The particle count was normalized to the cell count. The values are presented as the means ± SDs (n=3). *p<0.05 by one-way ANOVA with post-hoc Tukey’s post hoc test. **G, H.** Representative images of NTA of EVs secreted in the rescue experiments with serine supplementation. **I, J.** Knockdown of PHGDH and PSPH with each siRNA in HCT116 and A549 cell lines. **K.** NTA of EVs secreted from HCT116 and A549 cells after PHGDH and PSPH silencing. The values are presented as the means ± SDs (n=3). *p<0.05 by one-way ANOVA with Dunnett’s test. **L**. Viability of HCT116 and A549 cells after PHGDH and PSPH silencing. Cell viability was measured with a CCK-8 assay. The values are presented as the means ± SDs (n=3). *p<0.05 by one-way ANOVA with Dunnett’s test.

To determine whether PSAT1 affects known EV secretory pathways, we investigated the protein levels of TSG101 and ALIX in the ESCRT-dependent pathway, RAB27A, RAB27b, RAB11, and VAP-A related to membrane transport, and CAV1 in caveolae-dependent secretion (10). Western blotting analysis showed no obvious difference in these protein levels between PSAT1-knockdown and control samples, suggesting that PSAT1 manipulations at least did not affect known EV secretion-related protein levels (**Figure S4E**).

To further confirm the importance of the serine-ceramide synthesis pathway for EV secretion, we performed knockdown of PHGDH, which is upstream of PSAT1, and PSPH, which is downstream of PSAT1 in the serine synthesis pathway. The knockdown efficiency of the PHGDH and PSPH by siRNAs was confirmed in the HCT116 and A549 cell lines at the protein level **(Figures 3I and 3J)**. As expected, PHGDH silencing significantly decreased EV secretion in both cell lines, while the EV number was significantly decreased only in A549 cells after PSPH silencing (**Figure 3K)**. Cellular serine levels were also decreased in PHGDH and PSPH knockdown cells (**Figure S5A**). EV secretion and cellular serine levels were significantly correlated in PSAT1, PHGDH, and PSPH knockdown cells (**Figure S5B**). Cell proliferation was not changed after knockdown, although PSPH siRNA slightly reduced the cell number in A549 cells (**Figure 3L**). We also examined the effect of gene silencing related to the ceramide synthesis pathway on the EV secretion. When genes such as SPTLC1, KDSR, CERS6, and DEGS1 were inhibited by siRNAs (**Figure S5C**), these gene silencing significantly decreased EV secretion in both HCT116 and A549 cell lines (**Figure S5D**). Because ceramide is one of the important lipids for EV formation, these data suggested that PSAT1-mediated serine-ceramide synthesis pathway was involved in the EV production.

To obtain ancillary data showing the importance of the serine-ceramide synthesis pathway for EV secretion, we used inhibitors such as NCT-503, a PHGDH inhibitor (22–23), Fumonisin B1, a ceramide synthesis inhibitor (24), and GW4869 (3), a neutral sphingomyelinase inhibitor (**Figure S5E)**. Treatment with the inhibitors of serine-ceramide synthesis pathway decreased EV secretion in the HCT116 and A549 cell lines (**Figures S5F and S5G**). These data further confirmed that the serine-ceramide synthesis pathway contributes to EV secretion in cancer cells.

### Involvement of PSAT1 in EV biogenesis

We next examined whether PSAT1 affected EV biogenesis. For this purpose, we co-immunostained CD63 as an EV surface marker and PSAT1 in PSAT1-silenced HCT116 and A549 cells. Immunofluorescence staining of CD63 (green) and PSAT1 (red) showed accumulation of CD63 signals in the cytoplasm after PSAT1 silencing **(Figures 4A and 4B)**. Likewise, the accumulation of CD63 signals was clearly observed after miR-891b transfection **(Figure S6A**) and serine-ceramide pathway inhibitor treatments, particularly NCT-503 treatment (**Figure 4C**). These observations suggested that CD63 protein, the EV surface marker, might not be properly loaded onto EVs because inhibition of the serine synthesis pathway reduces the amounts of EVs production in cancer cells. To further elucidate the role of PSAT1 on EV biogenesis in detail, we co-immunostained CD63 with the early endosome marker EEA1 or the late endosome marker RAB7. As shown in **Figure S6B**, very little overlap between EEA1 and CD63 was observed in either NC or siPSAT1, while the majority of CD63 and RAB7 overlapped following NC control (**Figure S6C**). Introducing siRNA for PSAT1 caused increased CD63 single positive area and less colocalization of CD63 and RAB7 (**Figures 4D, 4E, and S6C**). This finding suggests that suppression of PSAT1 might be involved in EV biogenesis at stages of late endosome formation.

**Fig. 4.**
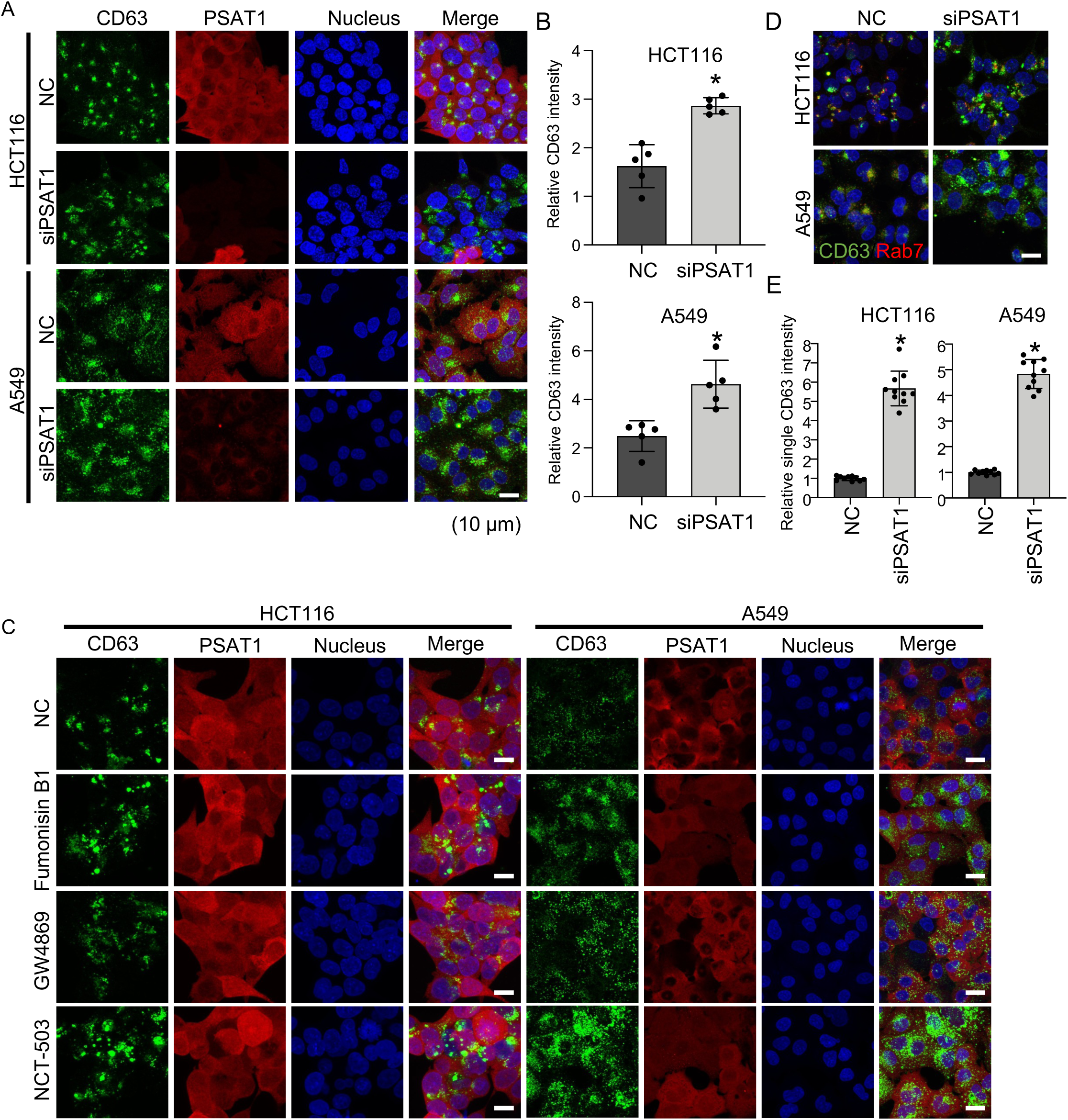
Investigation of the association between PSAT1 expression and EV biogenesis. **A.** Costaining of the CD63 (green) with PSAT1 (red) and nucleus (Hoechst 33442, blue) in PSAT1-silenced HCT116 and A549 cells. **B**. Quantification of the sum of the CD63-positive area in HCT116 and A549 cells in the PSAT1-silenced and control groups (n=5, *p<0.05 by Student’s t-test). **C.** Co-immunostaining of the CD63 (green) with PSAT1 (red) and nucleus (Hoechst 33442, blue) in serine-ceramide pathway inhibitor-treated HCT116 and A549 cells. **D.** Co-immunostaining of RAB7, a late endosome marker (red), with CD63 (green). **E.** Quantification of the sum of the CD63 single-positive area in HCT116 and A549 cells in the PSAT1-silenced and control groups (n=10, *p<0.05 by Student’s t-test). Scale bars; 10 μm.

### PSAT1 regulates EV secretion in multiple cancer cell lines

We sought to examine whether the PSAT1-controlled EV secretion mechanism is conserved in other cancer cells and normal cells. First, we evaluated the expression levels of PSAT1 in colorectal cancer cell lines (HCT15, COLO201, COLO205, and HT-29), normal colon fibroblasts (CCD-18co), lung cancer cell lines (A427, H1650, and H2228), and normal lung epithelial cells (HBEC). The expression levels of PSAT1 were generally higher in both types of cancer cells than in the corresponding normal cell lines **(Figure 5A)**. The PSAT1 expression level was positively correlated with EV secretion (**Figure 5B**). In contrast, the expression of miR-891b was higher in normal cells **(Figure S7A)**. Then, we analyzed PSAT1 expression in clinical samples deposited in the public database Oncomine. We confirmed the higher expression of PSAT1 in colon cancer, lung cancer, ovarian cancer, breast cancer, melanoma, head and neck cancer and multiple myeloma but not in pancreatic cancer (**Figure 5C)**. Furthermore, a high expression level of PSAT1 was associated with poor overall survival in lung cancer patients, particularly at the early stage. **(Figure S7B)**. In colorectal cancer, a previous study reported that a high expression level of PSAT1 was found to be related to poor prognosis (25). Additionally, a high expression level of PSAT1 was related to poor prognosis in many types of cancer, including breast cancer, pancreatic ductal adenocarcinoma, liver hepatocellular cancer, sarcoma, kidney renal clear cell carcinoma, and uterine corpus endometrial carcinoma. In head and neck cancer, the difference was not statistically significant but aligned in a similar trend with other cancers (**Figure S7C**). These data indicated that PSAT1 is associated with cancer progression in multiple types of cancers.

**Fig. 5.**
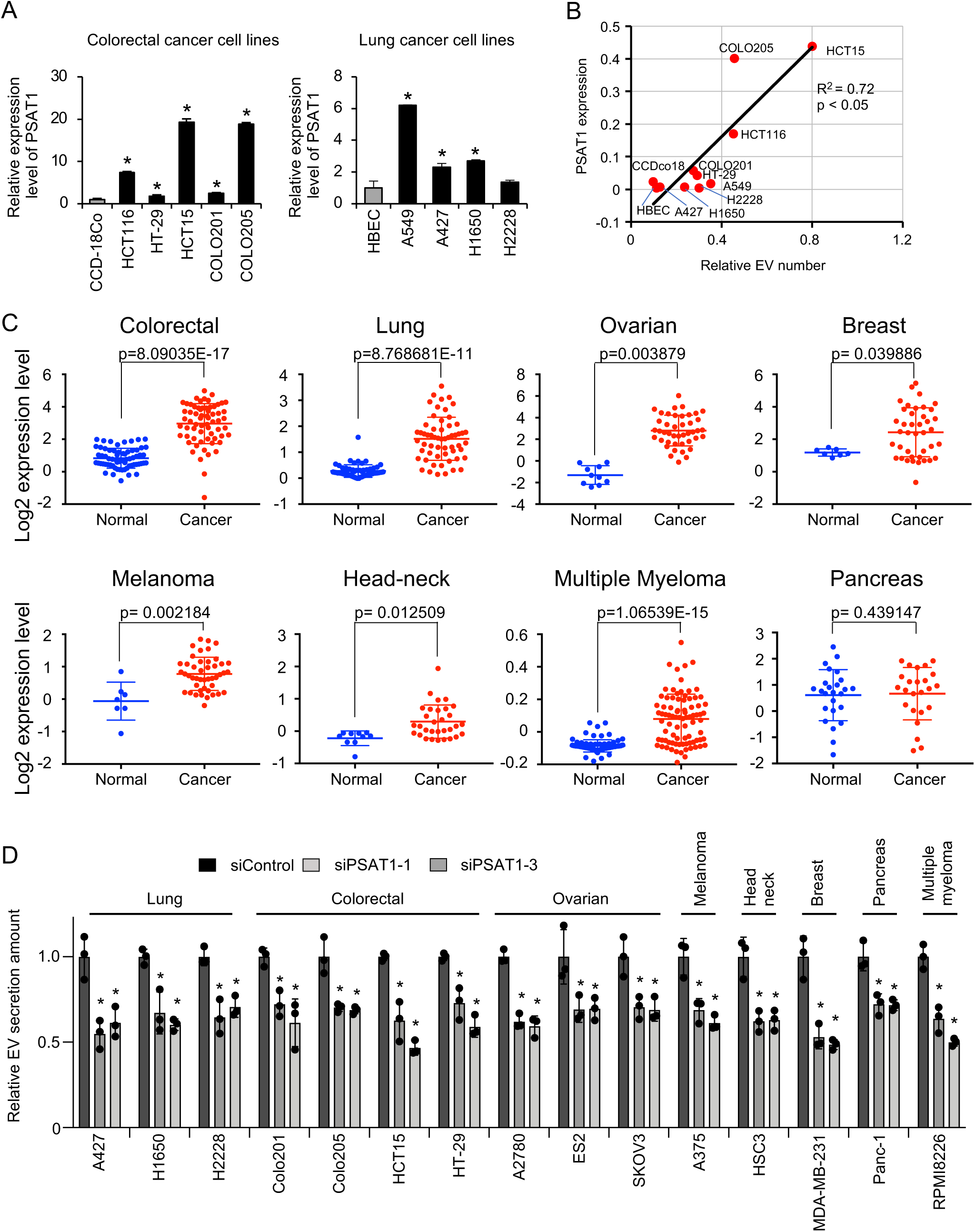
PSAT1 universally regulates EV secretion in cancer. **A.** qRT–PCR analysis of the expression level of PSAT1 across colorectal and lung cancer cell lines. Expression levels were normalized to β-actin expression. The values are presented as the means ± SDs (n=3). *p<0.05 by one-way ANOVA with Dunnett’s test. **B.** Positive correlation between PSAT1 expression and EV secretion across cell lines. **C.** Comparison of PSAT1 expression levels between cancer and normal tissues in several types of cancer based on the Oncomine database. p values were calculated by Student’s t test. **D.** The effect of siPSAT1 (siPSAT1-1 and siPSAT1-3) on EV secretion across cancer cell lines. EV secretion was measured by NTA. The values are presented as the means ± SDs (n=3). *p<0.05 by one-way ANOVA with Dunnett’s test in each cell line.

Next, we confirmed the contribution of PSAT1 to EV secretion in other cancer types. We cultured a variety of cancer cell lines, such as ovarian (A2780, ES2, and SKOV3), melanoma (A375), head and neck (HSC3), breast (MDA-MB-231), pancreatic (Panc-1), and multiple myeloma (RPMI8226) cell lines. EVs secreted from these cells were collected by ultracentrifugation after PSAT1 siRNA transfection (both siPSAT1-1 and siPSAT1-3). The EV particle number was significantly decreased in all cell lines after the PSAT1 silencing (**Figure 5D**). Although we did not observe a significant difference in PSAT1 expression between normal and cancer cells in pancreatic cancer **(Figure 5C, bottom right)**, the EV particle number in Panc-1 cells was decreased after PSAT1 silencing **(Figure 5D)**. These data suggested that PSAT1 positively regulates EV secretion in multiple types of cancer.

### Increased EV secretion by PSAT1 in breast cancer promoted osteoclast differentiation

During the analyses of PSAT1 expression in a number of cancer cell lines, we observed higher expression of PSAT1 in metastatic breast cancer cell lines than in their parental cell lines **(Figure 6A**). The bone metastatic cell line BM02 (**Figure 6A, top panels**) and lymph node metastatic cell line D3H2LN (**Figure 6A, bottom panels**) produced greater amounts of PSAT1 protein than their parental cell lines MCF7 and MDA-MB-231, respectively. We also found that metastatic cell lines secreted higher amounts of EVs than parental cells (**Figure 6B**). Next, we decided to investigate how PSAT1-controlled EV secretion was involved in tumor malignancy. For this purpose, we chose the MCF7 cell line and its bone metastatic derivative, BM02, and established cells with stable PSAT1 overexpression (**Figure 6C**). Overexpression of PSAT1 significantly increased the amount of EVs secreted from both MCF7 and BM02 cells (**Figures 6D, and S8A**). Transmission electron microscope (TEM) analysis also confirmed increased EV production in MVB of PSAT1-overexpressing MCF7 cells (**Figure S8E, left panel**). Because BM02 cells have bone metastatic ability, we investigated whether BM02 cell-derived EVs influence osteoclast differentiation in vitro. Using a Boyden chamber coculture system, osteoclast differentiation of RAW 264.7 cells, a macrophage cell line, was examined by TRAP (tartrate-resistant acid phosphatase) staining (**Figures 6E and 6F**). PSAT1-overexpressing MCF7 cells treated with soluble RANKL (sRANKL) treatment significantly induced osteoclast differentiation, while PSAT1-overexpressing BM02 showed a tendency to increase osteoclast differentiation. In contrast, when PSAT1 expression was suppressed in BM02 cells with shRNA (**Figures S8B**), EV secretion was inhibited (**Figures S8C, and S8D**). TEM analysis also confirmed decreased EV production in MVB of PSAT1 knockdown BM02 cells (**Figure S8E, right panel**). Then, in the co-culture with the PSAT1 knockdown BM02 cells, osteoclast differentiation in RAW 264.7 cells were inhibited (**Figures 6G and 6H**). Osteoclast differentiation was also stimulated by supplementation with PSAT1-overexpressing cells-derived EVs isolated by a conventional ultracentrifugation method (**Figures 6I and 6J**). We also investigated the effect of PSAT1 expression on normal cells. To this end, we overexpressed PSAT1 in CCD-18Co, normal colon fibroblasts and examined the effects on EV secretion and osteoclast differentiation. While EV secretion from PSAT1-overexpressing CCD-18Co cells was significantly increased (**Figures S9A-S9C**), osteoclast differentiation was not induced in RAW 264.7 cells cocultured with PSAT1-overexpressing CCD-18Co cells (**Figure S9D**). These data indicated that PSAT1 itself is not important for osteoclast differentiation, suggesting that PSAT1-induced cancer EVs are critical for osteoclast differentiation in a bone metastatic breast cancer cell line.

**Fig. 6.**
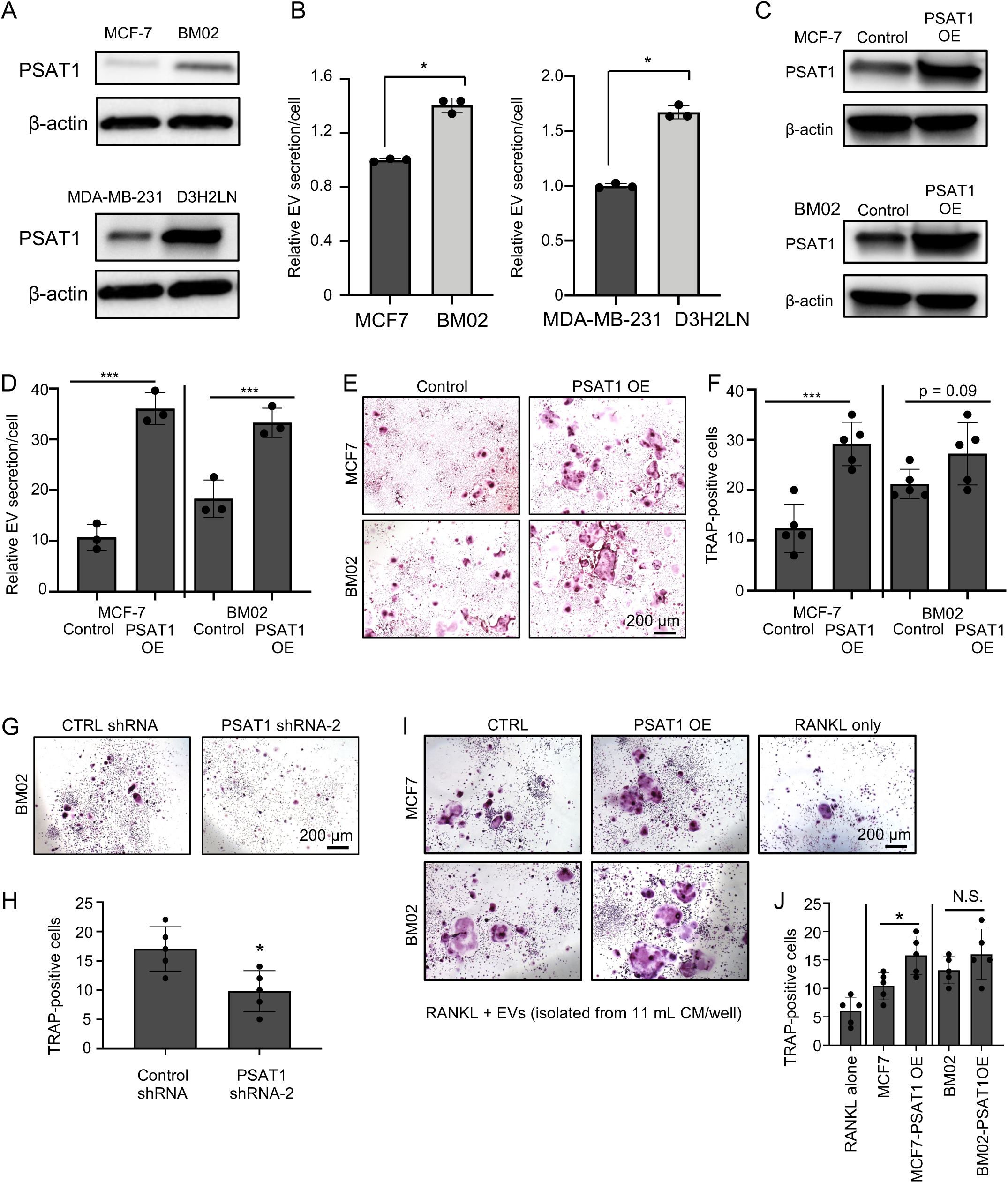
PSAT1-controlled EV secretion induces osteoclast differentiation in bone metastatic breast cancer cells. **A.** Western blot analysis of PSAT1 in parental cells (MCF7 and MDA-MB-231) and the corresponding metastatic cells (BM02, a bone-metastatic derivative of MCF7 cells; and D3H2LN, a lymph node-metastatic derivative of MDA-MB-231 cells). **B.** Relative EV secretion levels in parental and metastatic cell lines. The number of EVs secreted was normalized to the cell number. The values are presented as the means ± SDs (n=3). *p<0.05 by Student’s t test. **C.** Western blotting analysis of PSAT1 in MCF7 and BM02 cell lines with PSAT1 overexpression. **D**. EV particle number counts in MCF7 and BM02 cell lines with PSAT1 overexpression. The number of EVs secreted was normalized to the cell number. The values are presented as the means ± SDs (n=3). ***p<0.001 and **p<0.01 by Student’s t test. **E**. Images of TRAP staining in the coculture system with RAW 264.7 cells and MCF7/BM02 cells with PSAT1 overexpression. Scale bar: 200 μm. **F**. Quantification of TRAP-positive cells in a coculture system with MCF7/BM02 cells with PSAT1 overexpression. The values are presented as the means ± SDs (n=5). ***p< 0.001 by Student’s t test. **G**. Images of TRAP staining in the coculture system with RAW 264.7 cells and PSAT1-knockdown BM02 cells. Scale bar: 200 μm. **H**. Quantification of TRAP-positive RAW 264.7 cells in the coculture system with PSAT1-knockdown BM02 cells. The values are presented as the means ± SDs (n=5). *p<0.05 by Student’s t test. **I**. Images of TRAP staining upon supplementation with EVs derived from control MCF7, MCF7 PSAT1 OE, control BM02, and BM02 PSAT1 OE cells. Scale bar: 200 μm. **J**. Quantification of TRAP-positive cells under each condition. The values are presented as the means ± SDs (n=5). *p<0.05 and N.S.: not significant by Student t test.

In support of this idea, we investigated whether PSAT1 influences EV cargo. We performed small RNA-seq on EV samples isolated from PSAT1-overexpressing MCF7, control MCF7, PSAT1-knockdown BM02, and control BM02 cells. No obvious differences in EV miRNAs were not observed among the PSAT1 expression statuses (**Figure S10A and Table S2**). As RANKL is a strong inducer of osteoclast differentiation, we examined whether PSAT1 overexpression affects the protein level of RANKL. Western blot analysis confirmed that RANKL protein level was not changed by PSAT1 overexpression, although the RANK level was slightly altered (**Figure S10B, left panels**). We also confirmed that PSAT1 overexpression did not influence the expression of typical EV markers such as CD9, CD63, and FLOT-1, and the particle size of EVs (**Figures S10B, right panels and S10C**), indicating that the enhanced osteoclast differentiation of MCF7 and BM02 cells was due to the increased number of EVs secreted.

### Enhanced PSAT1 expression contributes to breast cancer metastasis via EV secretion

We sought to investigate whether PSAT1-controlled EV secretion influences bone metastasis using a bone metastatic breast cancer cell line. We utilized a bone metastasis model established by caudal artery injection of MCF7 and BM02 cells (26), and metastatic processes were monitored over time with an in vivo imaging system (IVIS). In vivo imaging revealed that PSAT1-overexpressing MCF7 cells significantly promoted bone metastasis (**Figures 7A and 7B**). Histological analysis indicated marked invasion of PSAT1-overexpressing MCF7 cells into the bone, accompanied by structural destruction (**Figure 7C**). Consistent with the increased osteoclast differentiation observed in vitro, increased osteoclast differentiation in bone from mice injected with PSAT1-overexpressing MCF7 cells was clearly observed by TRAP staining (**Figure 7D**). A similar trend was also observed in the model established with PSAT1-overexpressing BM02 cells (**Figures S11A-S11D**). Computed tomography clearly showed an abnormal bone structure in mice injected with PSAT1-overexpressing BM02 cells (**Figure S11E**). Likewise, we examined the effect of PSAT1-knockdown BM02 cells on bone metastasis. PSAT1 inhibition in BM02 cells significantly decreased bone metastasis (**Figures 7E and 7F**). Histological analysis indicated decreased invasion of PSAT1-knockdown BM02 cells into the bone (**Figure 7G**). Osteoclast differentiation, as evaluated by TRAP staining, showed a tendency to be decreased in bone from mice injected with PSAT1-knockdown BM02 cells (**Figure 7H**). Next, the effect of PSAT1-driven EVs on cancer cells themselves was examined. EVs were isolated from the same amount of conditioned medium from control MCF7 cells and PSAT1-overexpressing MCF7 cells by a conventional ultracentrifugation method. MCF7 cells were pretreated with the EVs isolated from these equal amounts of conditioned medium and then transplanted via caudal artery injection for evaluation of bone metastasis. In vivo imaging revealed that only a subtle increase in bone metastasis in mice injected with MCF7 cells pretreated with PSAT1-overexpressing MCF7 cell-derived EVs (**Figure S11F**), implying that the effect of EVs on the cancer microenvironment, rather than on the cancer cells themselves, was more pronounced. In addition to the bone metastasis model established with bone metastatic breast cancer cell lines, using a lung metastasis model with tail vein injection, we tested whether PSAT1-overexpressing breast cancer cells enhance lung metastatic potential. In vivo imaging revealed that PSAT1 overexpression in MDA-MB-231 cells significantly promoted lung metastasis (**Figures 7I and 7J**). Histological analysis of the lungs showed increased micrometastatic foci in mice injected with PSAT1-overexpressing MDA-MB-231 cells (**Figures 7K and 7L**). In brief, these data suggest that PSAT1 functions to universally enhance behaviors related to the malignant potential of cancer, such as metastasis, by increasing the secretion of EVs.

**Fig. 7.**
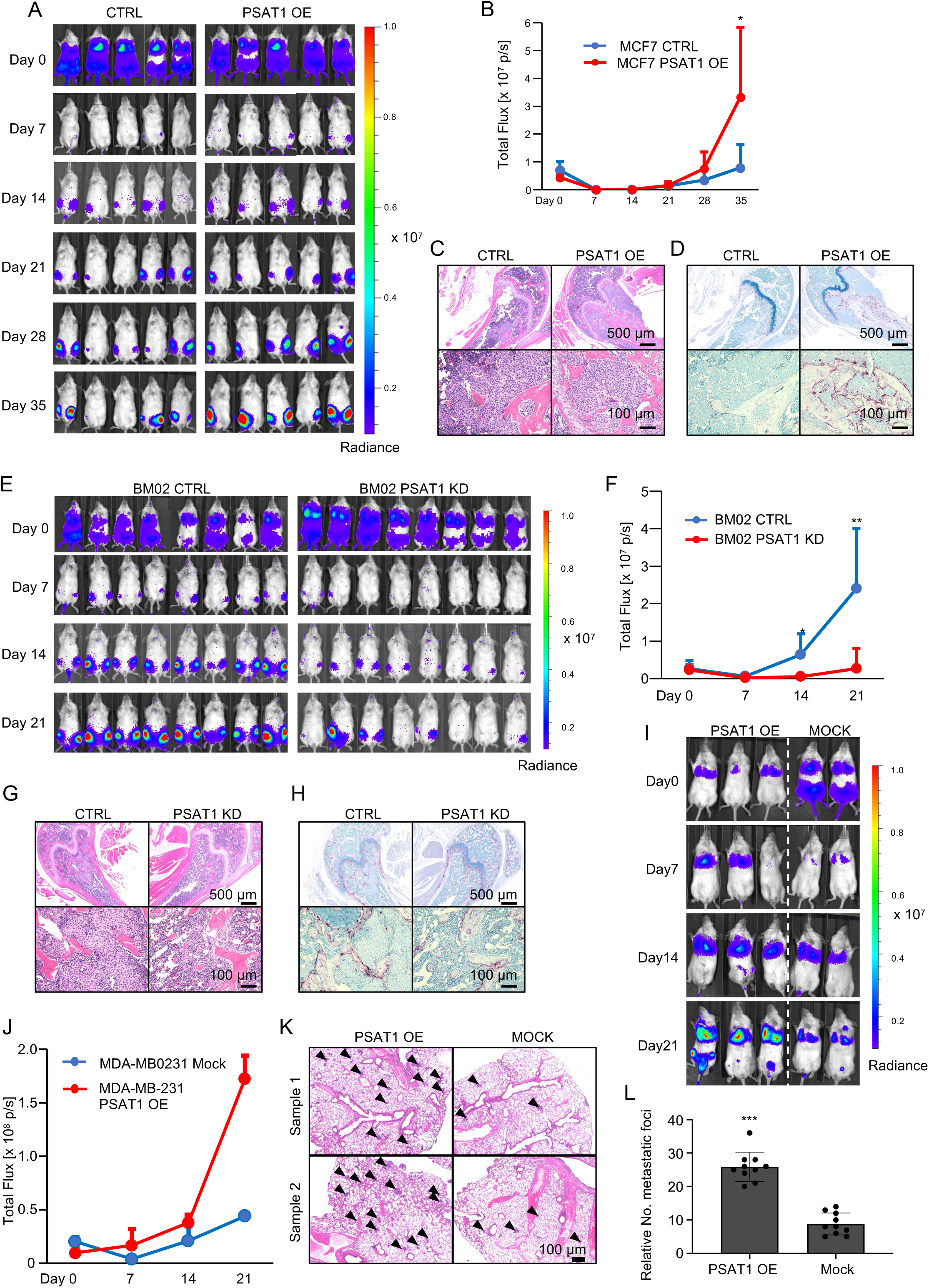
PSAT1 facilitates metastasis in metastatic breast cancer models. **A.** IVIS imaging of MCF7 PSAT1 OE cells. Images acquired on days 0, 7, 14, 21, 28, and 35 are shown. **B**. Photon counting of MCF7 PSAT1 OE cells in the bone metastasis mouse model. The values are presented as the means ± SDs (n=5). *p<0.05 by Student t test. **C**, **D**. HE staining of MCF7 PSAT1 OE cells (**C**). TRAP staining of MCF7 PSAT1 OE cells (**D**). Top panels: low magnification; bottom panels: high magnification. Scale bars: 500 μm for the top panels and 100 μm for the bottom panels. **E**. IVIS imaging of BM02 PSAT1 KD cells. Images acquired on days 0, 7, 14, and 21 are shown. **F**. Photon counting of BM02 PSAT1 KD cells in the bone metastasis mouse model. The values are presented the means ± SDs (n=5). *p < 0.05 and **p < 0.01 by Student t test. **G, H**. HE staining of BM02 PSAT1 KD cells (**G**). TRAP staining of BM02 PSAT1 KD cells (**H**). Top panels: low magnification and bottom panels: high magnification. Scale bars: 500 μm for the top panels and 100 μm for the bottom panels. **I**. IVIS imaging of MDA-MB-231 PSAT1 OE cells injected via tail vein. Images acquired on days 0, 7, 14, and 21 are shown. MDA-MB-231 PSAT1 OE, n=3; MDA-MB-231 mock, n = 2. **J**. Photon counting of MDA-MB-231 PSAT1 OE cells in the lung metastatic model. The values are presented as the means ± SDs. **p<0.01 by Student t test. **K.** HE staining of lung tissues from MDA-MB-231 PSAT1 OE cells. The arrowheads indicate metastatic foci in the lungs. Scale bars: 100 μm. **L**. Number of metastatic foci formed from MDA-MB-231 PSAT1 OE cells. The values are presented as the means ± SDs (n=8). ***p<0.001 by Student t test.

## Discussion

Cancer-derived EVs exacerbate cancer malignancy through modulating microenvironmental cells to tumor-supportive (27–28), stimulate invasion and metastasis (29–30), and contribute to drug resistance (6, 31–32). Therefore, intervening the transfer of cancer cell-derived EVs is expected to be a new therapeutic strategy for cancer (33–34). In this study, we identified miR-891b as a miRNA decreasing EV secretion from cancer cells by downregulating PSAT1, an enzyme catalyzes serine synthesis. In addition to silencing of the PSAT1, PSPH, and PHGDH genes in serine synthesis pathway and the SPTLC1, KDSR, CERS6, and DEGS1 genes in ceramide synthesis pathway, treatment with inhibitors such as NCT-503, Fumonisin B1, and GW4869 also decreased EV secretion. These data clearly indicated that the serine-ceramide pathway is important for EV secretion in cancer cells. Moreover, our findings suggest that the effect of PSAT1 on EV secretion is conserved across multiple types of cancers.

Our results showed that the increased secretion of EVs from cancer cells may explain the higher expression of PSAT1, i.e. aberrant serine metabolism, in cancer cells than in normal cells. Serine levels are correlated with the EV secretion under several conditions. Serine-related metabolic pathways have been reported to increase cancer cell proliferation (35–37). For example, HCT116 cells are highly dependent on de novo biosynthesis of serine and glycine (36, 38). Serine is also critical for the survival and growth of cancer cells and has been shown to play a central role in metabolic reprogramming. The serine metabolism pathway is important for synthesis of ceramide, which is one of the EV components (11). As for another biological aspect of PSAT1-mediated EV secretion, we have presented data on the effects of PSAT1 and the serine-ceramide synthesis pathway on EV biosynthesis in this study; however, we believe that more conclusive data can be obtained by further analysis such as live imaging techniques with ultra-high resolution microscopy. In addition to these, small-RNA sequencing showed that the components of EV did not change after PSAT1 modulation. Additionally, normal cells with PSAT1 overexpression secreted more EVs, but these EVs could not induce osteoclast differentiation. Our results suggested that activation of serine-related pathways may contribute to the increase of EVs release by supplying a membrane component of EVs in cancer cells.

In some cancer cells, the copy number of the PHGDH gene, which encodes the enzyme involved in the first step of serine synthesis, was found to be increased due to gene duplication (39–40). Increased PHGDH expression resulted in morphological abnormalities in mammary epithelial cells (40). In addition, the expression level of PHGDH affected breast cancer metastasis (41). Rossi M et al. revealed that breast cancer cells have a heterogeneous expression levels of PHGDH. Endothelial cells can induce PHDGH loss in cancer cells. Low PHGDH expression in these cells promotes cancer metastasis via aberrant protein glycosylation, including increased sialylation of integrin αvβ3. The migration and invasion of these cells are potentiated. In contrast, at the metastatic site, the expression level of PHGDH is restored, which increases the proliferation of these cells. Previous studies indicate that abnormal serine metabolism is linked with cancer development and tumor initiation. PSAT1 catalyzes 3-hydroxypyruvate to phosphoserine reaction and also converts glutamine to alpha-ketoglutarate (α-KG), one of the molecules consisting tricarboxylic acid cycle (TCA cycle). Hwang IY et al. showed that PSAT1 is involved in maintaining intracellular α-KG levels in both cancer and normal cells (42). In addition, PSAT1 is associated with vitamin B6 metabolism (43–44). In our screen, we identified the miR-891b target gene PDXK, which is associated with vitamin B6 metabolism. Silencing of PDXK with siRNA decreased EV secretion (**Figure S2C**). It is likely that vitamin B6 metabolism may also be involved in EV secretion with the serine synthesis pathway. We showed that miR-891b regulates EV secretion via multiple pathways because miRNAs posttranscriptionally regulate the expression of a large number of genes (45). Focusing not on a single pathway but on the whole network of processes becomes important when considering cancer progression.

This study has several limitations. First, the ExoScreen method used in the presented study can detect only CD63 or CD9 positive small EVs. We also used NTA system to monitor the EV particle size and number by NTA; however, the analysis largely showed a single peak so that the small EVs without the expression of these markers could not be differently evaluated. Although the molecular heterogeneity of EV populations is currently the focus of attention, the results in this study may not be universal to all EV populations, and it is possible that certain EV populations may be affected by the serine-ceramide pathway in cancer. Further analysis is needed to evaluate the relationship between CD63 or CD9-negative small EVs and the serine synthesis pathway. Second, to directly show an importance of serine on EV production, serine depletion from the culture medium is another strategy. In this study, we genetically or pharmacologically suppressed serine-ceramide pathway; however, due to the technical difficulty, we could not show the result of serine depletion, although we showed the results of serine addition into serine-free medium. Third, we could not draw conclusions regarding the importance of EVs in the cancer metastasis model. Although our in vitro and in vivo experimental data clearly indicated the effect of PSAT1 in the bone metastatic breast cancer model, we failed to show direct evidence of EV function in vivo by systematic injection of isolated EVs into mice in the bone metastasis model. Direct injection of a large amount of EVs via the tail vein is technically challenging because repeated tail vein injections of EVs damage the blood vessels in the tails. Additionally, the amount of EVs that could reach the metastatic site was uncertain, because we speculated that many EVs were trapped by endothelial cells in blood vessels.

In summary, our study revealed novel mechanisms of EV secretion via PSAT1-related serine synthesis in multiple types of cancer. In addition, our in vivo experiments revealed that PSAT1-controlled EV secretion might facilitate cancer metastasis by educating the cancer microenvironment. These findings may provide a novel cancer treatment strategy through targeting of serine synthesis and EV secretion.

## Materials and Methods

### Cell culture

HCT116, HCT15, COLO201, COLO205, CCD-18co, A549, NCI-H1650, NCI-H2228, A427, RPMI8226, A375, Panc-1, HSC3, HSC4, A2780, SKOV3, ES2, SCC25, MCF7, MDA-MB-231, and RAW264.7 were purchased from ATCC (American Type Culture Collection). Human bronchial epithelial cells (HBECs) were purchased from Lonza. The metastatic MDA-MB-231 cell line D3H2LN was purchased from Xenogen. Bone metastatic cells (BM02) were generated from MCF7 cells (46–47). Cells were cultured in Dulbecco’s modified Eagle’s medium (DMEM) and RPMI 1640 medium (Gibco) supplemented with 10% fetal bovine serum (FBS, Gibco) and 1% antibiotic-antimycotic (Gibco), and all cells were maintained in an atmosphere of 95% air and 5% CO_2_. Normal human epithelial cells were cultured in keratinocyte-SFM medium (Gibco) supplemented with 1% antibiotic-antimycotic (Gibco).

### EV isolation

Cancer cells were seeded in normal medium and cultured for 24 hours, and the medium was then replaced with serum-free advanced DMEM supplemented with 2 mM L-glutamine and 1% antibiotic-antimycotic (Gibco). After incubation for 48 hours, the conditioned medium was filtered with a 0.22 µm filter (SLGVR33RS, Millipore) and centrifuged at 2,000 × g for 10 min to remove cell debris. For EV isolation, the conditioned medium was ultracentrifuged at 110,000 × g for 70 min at 4 °C. The EV protein concentration was measured by using a Micro BCA Protein Assay Kit (23235, Thermo Fisher Scientific). The collection method, validation, and quantification methods suffice the MISEV2018 guidelines (48). Microvesicles were isolated from HCT116 and A549 cells by centrifugation. Conditioned medium was collected and centrifuged at 2,000 × g for 10 min to remove cells and cell debris. The supernatant was first filtered through a 0.8 µm filter (1-6893-02, Merck) and was then centrifuged at 10,000 × g for 40 minutes at 4 °C to pellet of microvesicles.

### Western blot analysis

The following antibodies were used as primary antibodies: mouse monoclonal anti-CD9 (clone 12A12, dilution 1:1000) and anti-CD63 (clone 8A12, dilution 1:1000) from Cosmo Bio. The anti-PSAT1 antibody (10501-1-AP, dilution 1:1000), anti-PHGDH antibody (14719-1-AP, dilution 1:1000) and anti-PSPH antibody (14513-1-AP, dilution 1:1000) were purchased from Proteintech Group. The anti-TSG101 antibody (612696, BD Biosciences, dilution 1:1000), anti-ALIX antibody (H00010015-B02, Abnova, dilution 1:1000), anti-RANK antibody (ab13918, Abcam, dilution 1:1000), anti-RANKL antibody (ab93718, Abcam, dilution 1:1000), anti-Flot-1 antibody (610820, BD Biosciences, dilution 1:1000), anti-RAB27A antibody (69295S, CST, dilution 1:1000), anti-RAB27B antibody (17572S, CST, dilution 1:1000), anti-RAB11 antibody (5589S, CST, dilution 1:1000), anti-CAV1 antibody (3267S, CST, dilution 1:1000), anti-VAP-A antibody (H00009218-M01, Abnova, dilution 1:1000), anti-SPTLC1 antibody (15376-1-AP, Proteintech, dilution 1:1000), anti-KDSR antibody (16228-1-AP, Proteintech, dilution 1:1000) anti-CER6 antibody (ab115539, Abcam, dilution 1:1000), and anti-DEGS1 antibody (ab185237, Abcam, dilution 1:1000) were used for Western blot analysis. Secondary antibodies (horseradish peroxidase-conjugated anti-mouse IgG, NA931, or horseradish peroxidase-conjugated anti-rabbit IgG, NA934; dilution 1:5000) were purchased from GE Healthcare. The same amount of protein was loaded in each lane of Mini-PROTEAN TGX gels (4–20%, Bio-Rad). The antibody signals were detected by chemiluminescence using ImmunoStar LD reagent (290-69904, FUJIFILM Wako), and analyzed with a luminescence imaging system (LAS-4000; Fujifilm Inc.).

### miRNA-based screening using the ExoScreen assay system

The following miRNA mimics and siRNAs were purchased from Ambion and were used for transient transfection, RAB27a siRNA (S11693), RAB27b siRNA (S11697), and nSMase2 siRNA (S30925). Allstar negative control siRNA (SI03650318) and TSG101 siRNA (SI02655184) were purchased from QIAGEN. The following antibodies were used for ExoScreen assay: mouse monoclonal anti-human CD9 (clone 12A12, Cosmo Bio) and anti-human CD63 (clone 8A12, Cosmo Bio). These antibodies were used to modify either acceptor beads or biotin following the manufacturer’s protocol. miRNA-based screening was performed as described previously (14). Cancer cells (5 × 10^3^ cells) were seeded in 96-well plates in DMEM supplemented with 10% FBS (Gibco) and 1% antibiotic-antimycotic (Gibco). After 24 hours of incubation, the AccuTargetTM Human miRNA Mimic Library, which was constructed based on miRBase ver. 21 (CosmoBio), was transfected into cells with DharmaFECT 1 Transfection Reagent (Dharmacon). The transfected cells were cultured in serum-free advanced DMEM. After 48 hours, the ExoScreen assay (21) was performed. The wells of a 96-well half-area white plate were filled with 10 μL of each sample and 15 μL of universal buffer mixture (5 nM biotinylated antibodies and 50 μg/ml AlphaLISA acceptor bead-conjugated antibodies). Combinations of “CD9 and CD9”, and “CD63 and CD63” were used. The plate was incubated for 1 hour at room temperature. Without a washing step, 25 μL of 80 μg/ml AlphaScreen streptavidin-coated donor beads were added. The plate was incubated in the dark for 30 minutes. The plate was read on an EnSpire Alpha 2300 Multilabel Plate Reader (PerkinElmer) with an excitation wavelength of 680 nm and emission wavelength of 615 nm. Background signals measured from filtered advanced RPMI medium or PBS were subtracted from the measured signals.

### Cell proliferation assay

Cell viability was determined using a Cell Counting Kit-8 (CK04, Dojindo) according to the manufacturer’s instructions, and the absorbance at 450 nm was measured using an EnVision Multilabel Plate Reader (PerkinElmer).

### Nanoparticle tracking analysis

The following miRNA mimics and siRNAs were purchased from Ambion and Dharmacon were used for transient transfection: PSAT1 siRNA (siGENOME SMARTpool siRNA M-010398, individual: siPSAT1 #1 D-010398-21-0002, siPSAT1 #2 D-010398-23-0002, siPSAT1 #3 D-010398-23-0002, siPSAT1 #4 D-010398-24-0002), SPTLC1 siRNA (siGENOME siSPTLC1 #1 D-006673-21, siSPTLC1 #2 D-006673-22), KDSR siRNA (siGENOME siKDSR #1 D-009936-04, siKDSR #2 D-009936-06), CER6 siRNA (siGENOME siCER6 #1 D-032207-01, siCER6 #2 D-032207-02), DEGS1 siRNA (siGENOME siDEGS1 #1 D-006675-01, siDEGS1 #2 D-006675-04), PHGDH siRNA (siGENOME SMARTpool siRNA M-009518) and PSPH siRNA (siGENOME SMARTpool siRNA M-011888). Cancer cells were seeded in normal medium and cultured for 24 hours. Then, the siRNAs were transfected with DharmaFECT 1 Transfection Reagent (Dharmacon) and after 24-hour the medium was replaced with serum-free advanced DMEM supplemented with 2 mM L-glutamine and 1% antibiotic-antimycotic (Gibco). After incubation for 48 hours, the conditioned medium was filtered with a 0.22 µm filter (SLGVR33RS, Millipore) and centrifuged at 2,000 × g for 10 min to remove cell debris. EVs were resuspended in PBS and further diluted for analysis with a NanoSight LM10-HS system (Quantum Design) according to the manufacturer’s protocol.

### Microarray analysis

A microarray analysis was performed as described previously (10). Agilent-072363 SurePrint G3 Human GE v3 8×60K Microarray (Agilent Technologies) was used. HCT116 cells and A549 cells were transfected with miRNA mimics, i.e., the miR-891b mimic (4464066, Ambion), and miRNA Mimic Negative Control #1 (4464058, Ambion). Total RNA was extracted from cultured cells using QIAzol reagent and a miRNeasy Mini Kit (217004, QIAGEN). RNA quantity and quality were determined using a NanoDrop ND-1000 spectrophotometer (Thermo Fisher Scientific) and an Agilent Bioanalyzer (Agilent Technologies).

### Quantitative real-time polymerase chain reaction

Total RNA was extracted from cultured cells using QIAzol reagent and a miRNeasy Mini Kit (217004, QIAGEN). RNA quantity and quality were determined using a NanoDrop ND-1000 spectrophotometer (Thermo Fisher Scientific). For qRT–PCR analysis, complementary DNA was generated from total RNA using a High Capacity cDNA Reverse Transcription Kit (4368814, Applied Biosystems). Total RNA was reverse transcribed using a TaqMan miRNA Reverse Transcription Kit from Applied Biosystems for the measurement of miR-891b expression (4366597, Applied Biosystems). The expression levels of mature miRNAs were measured using real[time qPCR with TaqMan Universal PCR Master Mix (4324018, Applied Biosystems). The data were collected and analyzed using StepOnePlus (Applied Biosciences). All quantitative mRNA expression data were normalized to that of β-actin. TaqMan probes for PSAT1 (Hs01107691), β-actin (Hs99999903), miR-891b (002210), and U6 (001093) were purchased from Applied Biosystems (PN4427975).

### Comprehensive miRNA sequencing in EVs

According to previous papers (49–50), comprehensive miRNA sequencing was performed. Sample EVs were isolated from MCF-7 cells, MCF-7 PSAT1 OE cells, BM02 cells and BM02 PSAT1 KD cells. RNA extraction was performed by using QIAzol reagent and a miRNeasy Mini Kit (217004, QIAGEN).

### Luciferase reporter assay

A fragment from the 3’UTR of PSAT1 containing the predicted target sequences of miR-891b was generated. The following annealed oligos, including the predicted target sequences of miR-891b (Thermo Fisher Scientific), were used to construct 3’UTR reporter vectors. PSAT1 3’UTR Sense: 5’-TCGAGAGTTAGATTTCAAACTTGCCTGTGGACTTAATAATGCAAGTTGCGATTAATTATTTCTGGAGTCATGGGAACACGC-3’; PSAT1 3’UTR Antisense: 5’-GGCCGCGTGTTCCCATGACTCCAGAAATAATTAATCGCAACTTGCATTATTAAGTCCACAGGCAAGTTTGAAATCTAACTC-3’; PSAT1 mut 3’UTR Sesne: 5’-TCGAGAGTTAGATTTCAAACTTGCCTGTGGACTTAATAATGCTTCAACGGATTAATTATTTCTGGAGTCATGGGAACACGC-3’; and PSAT1 mut 3’UTR AS: 5’-GGCCGCGTGTTCCCATGACTCCAGAAATAATTAATCCGTTGAAGCATTATTAAGTCCACAGGCAAGTTTGAAATCTAACTC-3’. The annealed oligos were ligated into the Xho I and Not I sites in the 3’UTR of the Renilla luciferase gene in the psiCHECK-2 plasmid (C8021, Promega). Each plasmid was transfected individually into HEK 293 cells with miR-891b using DharmaFECT 1 Transfection Reagent (T-2001-03, Dharmacon). Cells were cultured for 48 hours after transfection; then, firefly luciferase activity was measured and normalized to Renilla luciferase activity. Dual-Glo (E2920, Promega) was used for the luciferase assay and, the absorbance at 450 nm was measured using an EnVision Multilabel Plate Reader (PerkinElmer).

### Immunofluorescence

Cancer cells (1 × 10^5^) were seeded into 35 mm uncoated glass dishes (3910-135-MYP, IWAKI) and 4-well glass chamber slides (154526, Thermo Fisher Scientific). After 24 hours of incubation, siPSAT1 and Allstar negative control siRNA were transfected into each cell line, and after another 24 hours of incubation, the medium was changed to serum-free advanced DMEM. Confocal microscopy was performed with an Olympus FV10i laser scanning microscope (Olympus). The primary antibodies used as follows: anti-CD63 antibody (clone 8A12, Cosmo Bio, dilution 1:200), anti-PSAT1 antibody (Proteintech. dilution 1:200), anti-EEA1 antibody (ab109110, Abcam, dilution 1:200) and anti-RAB7 antibody (ab137029, Abcam, dilution 1:200)). Secondary antibodies conjugated to the following fluorophores were used: Alexa Fluor 488 (Thermo Fisher Scientific) for PSAT1, EEA1, and RAB7; Alexa Fluor 594 for CD63. Counter staining was performed with Hoechst 33442(Thermo Fisher Scientific).

### Serine supplementation experiment

MEM (Gibco) supplemented with 1 × insulin, transferrin, selenium solution (ITS-G, 100X, 41400045, Thermo Fisher Scientific), and 1 × MEM vitamin solution (100X, 11120052, Thermo Fisher Scientific), with or without serine (4 mM) and with or without ceramide (0.5 μM), was used. Serine was purchased from Sigma–Aldrich (S5386) for the serine supplementation experiment. Ceramide was purchased from Cayman Chemical (10681) for the ceramide supplementation experiment. Cancer cells were seeded into 96-well plates (5 × 10^3^ cells) or 6-well plates (1.5 × 10^5^ cells) and were then incubated for 24 hours. Then, siPSAT1 and Allstar negative control siRNA were transfected into each cell line. After 24 hours of incubation, the medium was changed to serum-free modified MEM. After another 48-hour incubation, the 96-well plates were used for the ExoScreen assay, and conditioned medium was collected from the 6-well plates. EVs were isolated from the conditioned medium, and NTA was then conducted. The inhibition rate of EV secretion was calculated based on the ExoScreen assay data and was normalized to the cell proliferation assay data.

### Inhibition of the serine synthesis pathway

The following reagents used for the experiments were purchased from the indicated companies: NCT-503 (S8619, Selleck), Fumonisin B1 (063-05873, FUJIFILM Wako) and GW4869 (S7609, Selleck). Cancer cells were seeded into 96-well plates (5 × 10^3^ cells) and were then incubated for 24 hours. Serine-related pathway inhibitors (NCT-503: 2.5 μM, Fumonisin B1: 1 μM and GW4869: 5 μM) were used at the indicated concentrations. After 24-hour incubation, the medium was changed to serum-free medium. After another 48-hour incubation, the 96-well plates were used for the ExoScreen assay.

### Measurement of serine concentration

Serine concentrations were measured by a serine kit DL-Serine Assay Kit (ab241027, Abcam) according to the manufacturer’s protocol. Cells were seeded into 6-well plates, and after 24-hour incubation, transfected with siRNAs. After another 48-hour incubation, cells were homogenized on ice with 100 μL of the supplied buffer, and cell lysates were collected from the 6-well plates.

### Public database analysis

Candidate target genes of miR-891b were predicted with the TargetScan database (http://www.targetscan.org/cgi-bin/targetscan/vert_72/targetscan.cgi?species=Human& gid=&mir_sc=&mir_c=&mir_nc=&mir_vnc=&mirg=mir-891b). PSAT1 expression data from cancer tissues and normal tissues deposited in the Oncomine database (https://www.oncomine.org/resource/login.html) were reanalyzed. Overall survival curves were calculated from the Kaplan–Meier Plotter database (https://kmplot.com/analysis/index.php?p=service&cancer=lung).

### Osteoclast differentiation & TRAP staining

RAW264.7 murine macrophages were cultured in minimum essential medium alpha (MEM-α) (Gibco) containing 10% FBS (Gibco) and 1% antibiotic-antimycotic (Gibco) and were maintained in an atmosphere of 95% air and 5% CO_2_. For osteoclast differentiation, RAW264.7 cells were cultured in the lower chamber for 120 hours with or without cancer cells in upper chamber in the presence of 10 ng/mL sRANKL (47187000, ORIENTAL YEAST CO. LTD.). Osteoclast differentiation was evaluated with a Tartrate-Resistant Acid Phosphatase (TRAP) Assay Kit (AK04F, Cosmo Bio). Stained cells with three or more nuclei were counted as TRAP-positive cells.

### Establishment of gene modified cell lines

#### a. Generation of PSAT1 OE cells

MCF7, BM02, and MDA-MB-231 cells (1 × 10^5^ cells) were seeded into 24-well dishes, and the PSAT1 overexpression vector (RC202475, Origene) was transfected by using Lipofectamine LTX Plus reagent (15338-100, Thermo Fisher Scientific). After 24 hours, the cells were transferred to 150 cm^2^ dishes containing 500 ng/mL geneticin (10131-027, Gibco). After geneticin selection, the cells that grew were established as PSAT1-overexpressing cells. The protein level of PSAT1 was validated by Western blotting.

#### b. Generation of PSAT1 KD cells and miR-891 OE cells

miR-891b overexpression vectors and shPSAT1 vectors were purchased from VectorBuilder (pLV[shRNA]-Puro-U6>hPSAT1[shRNA#1-#3] (VB900137-9279mqv, VB900137-9280eeg, VB900137-9281qvt), pLV[miRNA]-EGFP:T2A:Puro-U6>hmiR-891b (VB221007-1091txu), pLV[Exp]-EGFP/Puro-EF1A>ORF_stuffer (VB010000-9389rbj)). PSAT1 knockdown vectors were cotransfected with the pLP1, pLP2, and VSVG plasmids into 293T cells; after 48 hours of culture, conditioned medium containing virus particles was collected and used to infect BM02 cells. The PSAT1 knockdown efficiency was determined by Western blotting. miR-891 overexpressing cells were also established by this method. The expression level of miR-891b was measured by qRT-PCR.

#### c. Generation of PSAT1 knockout (KO) cells

The PSAT1 KO A549 cell line was established by CRISPR-Cas9 gene editing. The PSAT1 KO vector was used (50). A549 cells were seeded in 24 well plates. The PSAT1 KO vector was transfected with Lipofectamine 3000 (L3000015, Thermo Fisher Scientific). Puromycin selection (1 μg/mL) was initiated after 48 hours of culture. Single cells were isolated by limiting dilution, and the KO sequences were confirmed by Sanger sequencing. The PSAT1 KO efficiency was confirmed by Western blotting. The KO vector was constructed according to a previous protocol (51). F: CACCgCAATACAGAGAATCTTGTGC; R: AAACGCACAAGATTCTCTGTATTGc

### In vivo experiments

Female NOD.CB-17-Prkdc <scid>/J mice (NOD/scid, Charles River Japan, Inc.) were used for the animal study. The experiments were approved by NCC (T22-001) and carried out in accordance with the “Guideline for Animal Experiments in National Cancer Center” and the 3Rs stipulated by the Animal Welfare Management Act. After anesthetization of mice with 2.5% isoflurane gas (114-13340-3, Pfizer), PSAT1-overexpressing MCF7 cells or PSAT1 knockdown BM02 cells (5 x 10^5^ cells/100 μL in PBS (-) solution) were injected through the caudal artery. A 30 mg/mL D-luciferin (LUCK-1G, GOLDBIO) solution was prepared. Bone metastasis was monitored every week by bioluminescence imaging using an IVIS imaging system. Before the tumor burden was increased to > 10% of the mouse body weight, the mice were dissected and tumor were harvested at the experimental endpoint. PSAT1-overexpressing MDA-MB-231 cells were injected through the caudal vein. Lung metastasis was monitored every week by bioluminescence imaging using the IVIS imaging system. In vivo images were analyzed with Living image software ver. 4.7.2. The number of metastatic foci was counted from HE-staining images. Computed tomography of bones in mice injected with PSAT1-overexpressing BM02 and control BM02 cells was performed with the R_mCT2 instrument (RIGAKU). After tissue collection (femur and lung), tissue embedding was performed at Koto Microbiology Laboratory (Japan), the tissue sections were prepared, and HE staining was performed. According to the manufacturer’s protocol (Tartrate-Resistant Acid Phosphatase Assay Kit, AK04F, Cosmo Bio), TRAP staining of tissue was performed on thin section.

### Transmission electron microscopy (TEM)

Cultured MCF7, MCF7 PSAT1 OE, BM02, and BM02 PSAT1 KD cells were washed with PBS and were fixed with 2% glutaraldehyde at 4 °C, followed by 2% osmium tetroxide for 3 h. Fixed cells were dehydrated through an ethanol series (50%, 70%, 80%, 90%, 100%, 100%, and 100% for 5 min each at RT) and propylene oxide, and embedded in EPON 812 (TAAB Laboratories Equipment Ltd., Aldermaston, UK.). Ultrathin sections (70–80 nm thick) were cut on an EMUC7i ultramicrotome (Leica Microsystems, Tokyo, Japan), stained with 2% uranyl acetate and lead stain solution. TEM was performed using a JEM-1400 Plus transmission electron microscope (JEOL Ltd., Tokyo, Japan) at an accelerating voltage of 100 kV.

### Data availability

RNA-seq data of the miR-891b transfected cell lines generated for this study are included within this article and in the supporting information. The GEO accession number for our RNA-seq data is GSE185312. Small RNA-seq data for the EVs derived from MCF7 and BM02 cells are shown in Table S2.

### Statistical analysis

The data presented in the bar graphs are the means ± SDs of at least three independent experiments. Differences between two groups were analyzed by Student’s t test. When multiple comparisons testing was required, the significance of differences in the mean values was determined using one-way ANOVA with Dunnett’s test or Tukey’s honestly significant difference (HSD) post hoc test. Differences with p<0.05 were considered to be statistically significant. The log-rank test was performed for survival analysis using the ‘survminer’ package in R version 3.6.1. Correlation coefficients were calculated by Spearman correlation analysis. The heatmap was created by using Morpheus software (https://software.broadinstitute.org/morpheus/).

## Author contributions

TY, JN and FU designed the study. AY and MKi were performed Comprehensive small RNA sequencing in EVs. TY, KI, and JN performed in vitro and in vivo experiments. TY, YY, MKu and YH analyzed data. TY, JN, YY, NNA and TO wrote the manuscript. All authors read and approved the final manuscript.

## Acknowledgement

We thank Ms. M. Abe for experimental support. Department of Extracellular Vesicle Science, Industrial-Academic Collaboration, Tokyo Medical University was supported by Theoria Science, Inc. Japan. We thank the Division for Medical Research Engineering, Nagoya University Graduate School of Medicine, for equipment usage or technical support. This research was supported by AMED under Grant Number JP18ae0101011; JP22ama221405; MEXT KAKENHI (Grant-in-Aid for Scientific Research (B): 21H02721; Grant-in-Aid for Challenging Research (Pioneering): 23K17397; Princess Takamatsu Cancer Research Fund (22–25436); Kobayashi Foundation for Cancer Research; Mishima Kaiun Memorial Foundation; and Foundation for Promotion of Cancer Research.

## Supplementary Information

**Table S1. The 82 overlapping genes in the Venn diagram as shown in Figure 2A. Table S2. Read counts obtained by small RNA-seq of MCF7 and BM02 cells with PSAT1 manipulations**

**Figure S1. Screening of miRNAs in HCT116 and A549 cells**. **A.** EV secretion from cancer cells with knockdown of EV secretion-related genes. Left: HCT116, Right: A549. Normalized to NC, negative control; showing the means ± SDs (n=3). *p<0.05 by one-way ANOVA with Dunnett’s test. **B.** Second screen of selected miRNAs in HCT116 and A549 cells. EV secretion was measured by the ExoScreen method. Signals were normalized to the CCK-8 cell viability assay data. **C.** The effect of the miR-891b mimic on EV secretion was evaluated in conditioned medium (CM) and purified EVs. The amount of secreted EVs per cell was evaluated by the signal intensity measured in the ExoScreen assay. The values are presented as the means ± SDs (n=3). *p<0.05 by Student’s t test. **D.** NTA in miR-891b-transfected HCT116 and A549. The particle count was normalized to the cell count. The values are presented as the means ± SDs (n=3). *p<0.05 by Student’s t test**. E.** Representative images of NTA in miR-891b-transfected HCT116 and A549. **F.** Kaplan–Meier survival curves based on miR-891b expression in rectal cancer and lung cancer (https://kmplot.com/analysis/).

**Figure S2. The effect of PSAT1 on EV secretion. A.** The effects of transfection of different siPSAT1 sequences of in HCT116 and A549 cancer cell lines. Knockdown efficiency of each siPSAT1 sequence in HCT116 and A549. **B**. Particle count data of each siPSAT1 sequence in A549 cells. The values are presented as the means ± SDs (n=3). *p< 0.05 by one-way ANOVA with Dunnett’s test. **C**. PDXK, RAB31, and VPS37B were downregulated by transfection of the corresponding siRNAs (final concentration: 10 nM). The EV secretion level was evaluated by the ExoScreen method. **D.** NTA was used to determine the EV amount after PDXK, RAB31, and VPS37B siRNA transfection. Signals were normalized according to the corresponding cell counts. **E**. The effect of PSAT1 overexpression in the PSAT1-knockout A549 cell line. Quantification of EV particle numbers under each condition. The values are presented as the means ± SDs (n=3). *p<0.05 and N.S.: not significant by one-way ANOVA with Tukey’s post hoc test. **F**. Western blot analysis of PSAT1 levels under each condition. **G**. Serine levels after PSAT1 overexpression in the PSAT1-knockout A549 cell line. *p<0.05 by one-way ANOVA with Tukey’s post hoc test. **H**. The effect of PSAT1 overexpression in miR-891b-manipulated cancer cell lines. EV particle numbers were counted by NTA. The values are presented as the means ± SDs (n=3). * p < 0.05 by one-way ANOVA with Dunnett’s test. **I**. Representative images of NTA of EVs in miR-891b- and PSAT1-manipulated cancer cell lines.

**Figure S3. The effect of PSAT1 manipulation on microvesicle secretion. A.** Representative images of NTA of microvesicles isolated from siPSAT1-transfected HCT116 and A549 cells. **B.** Quantification of microvesicles particle numbers after PSAT1 knockdown in HCT116 and A549 cell lines. The particle numbers were counted by NTA. The values are presented as the means ± SDs (n=3). N.S.: not significant by Student’s t test.

**Figure S4. The effect of the serine-ceramide pathway on EV secretion. A.** Evaluation of the cell growth in medium with/without serine under PSAT1 manipulations. The growth of HCT116 and A549 cells was measured on days 0, 1, and 2 by a CCK-8 assay. The expression of PSAT1 was suppressed with siRNA, and cell growth was monitored in medium with/without serine (400 μM). The values are presented as the means ± SDs (n=3). **p<0.01 and ***p<0.001 by Student’s t test. **B**. representative NTA images of EV particles with/without serine addition. 4 mM serine was added into each. **C**. NTA of EVs secreted from HCT116, A549, MCF7, and BM02 cells with/without serine. The values are presented as the means ± SDs (n=3). *p<0.05 by Student’s t-test. **D.** Effect of ceramide on EV secretion under PSAT1 manipulation. Ceramide (0.5 μM) was added to HCT116 and A549 cells with/without siPSAT1 transfection. The particle number was determined by NTA. The values are presented as the means ± SDs (n=3). *p<0.05 by one-way ANOVA with Tukey’s post hoc test. **E**. Western blot analysis of proteins related to EV secretion. TSG101 and ALIX: ESCRT-dependent pathway, RAB27A, RAB27b, RAB11, VAP-A: membrane trafficking, and CAV1: Caveola-dependent secretion. PSAT1 expression was transiently inhibited by siPSAT1 in A549 and HCT116 cancer cell lines. In the BM02 cell line, PSAT1 was stably inhibited via shPSAT1. β-Actin was used as a loading control.

**Figure S5. The effect of gene silencing in the serine-ceramide pathway on EV secretion. A**. Measurement of serine levels under PSAT1 manipulations. After knockdown of PSAT1, PSPH, and PHGDH, total serine levels were measured based on fluorescence in the HCT116 and A549 cancer cell lines. The values are presented as the means ± SDs (n=3). *p<0.05 by one-way ANOVA with Dunnett’s test. **B**. Correlations between serine relative serine levels and relative EV secretion in HCT116 and A549 cancer cells with PSAT1, PSPH, and PHGDH knockdown. **C.** Confirmation of knockdown of ceramide synthesis pathway genes by western blot. β-actin was used as a loading control. **D.** Effect of gene silencing of the ceramide synthesis pathway on EV secretion. Particle number counting of EVs with siRNAs in HCT116 and A549 cell lines. The particle number was determined by NTA. The value in NC samples is set to 1.0. The values are presented as the means ± SDs (n=3). *p<0.05 by one-way ANOVA with Dunnett’s test. **E**. Serine-ceramide pathway and its inhibitors. NCT-503 is a PHGDH inhibitor, Fumonisin B1 is a ceramide synthesis inhibitor, and GW4869 is a neutral sphingomyelinase inhibitor. **F.** Effect of inhibitors of the serine synthesis pathway on EV secretion. Particle number counting of EVs with inhibitors in HCT116 and A549 cell lines. The particle number was determined by NTA. The value in NC samples is set to 1.0. The values are presented as the means ± SDs (n=3). *p<0.05 by one-way ANOVA with Dunnett’s test. **G.** Representative images of NTA of EVs from serine synthesis pathway inhibitor-treated HCT16 and A549 cells.

**Figure S6. The effects of miR-891b or PSAT1 silencing on EV biogenesis. A.** Co-immunostaining of CD63 (green) and PSAT1 (red) with the nucleus (Hoechst 33442, blue) in miR-891b-transfected cells. **B.** Co-immunostaining of EEA1, an early endosome marker (red), with CD63 (green). **C.** Co-immunostaining of RAB7, a late endosome marker (red), with CD63 (green).

**Figure S7. Kaplan–Meier plot based on PSAT1 expression across cancer types. A.** qRT-PCR analysis of miR-891b expression in normal cells. the expression levels of miR-891b in CCD-18Co cells and HBECs were measured, and the miR-891b expression levels were normalized to the U6 expression levels. *p<0.05 by Student’s t test. **B.** Kaplan–Meier survival curve based on PSAT1 expression in lung cancer (all stages, stage I, II and III). **C**. Kaplan–Meier survival curves of 8 cancer types.

**Figure S8. Characterization of PSAT1 overexpression and knockdown in breast cancer cells. A.** Representative NTA images of EVs from PSAT1 overexpressed MCF7 and BM02 cells. **B.** qRT-PCR analysis of PSAT1 expression in the BM02 cell line with PSAT1 knockdown. *p<0.05 by one-way ANOVA with Dunnett’s test. **C.** Representative NTA images of EVs from PSAT1 knockdown BM02 cells. **D.** EV particle numbers count in the BM02 cell line with PSAT1 knockdown. The values are presented as the means ± SDs (n=3). *p<0.05 by one-way ANOVA with Dunnett’s test. **E.** TEM images of PSAT1 overexpressed MCF7 and PSAT1 knockdown BM02 cells. Scale bars; 200 nm for upper panels and 2 μm for lower panels.

**Figure S9. The effect of PSAT1 overexpression on EV secretion in normal cells. A-C.** PSAT1 induced EV secretion in CCD-18Co cells. PSAT1 overexpression in CCD-18Co cells was confirmed by western blotting (**A**). EV secretion in CCD-18Co PSAT1 OE cells (**B**). The particle number was determined by NTA. The values are presented as the means ± SDs (n=3). **p<0.05 by Student’s t test. Representative NTA images of CCD-18Co PSAT1 OE cells (**C**). **D.** TRAP staining of osteoclast differentiation in the coculture with CCD-18Co cells, CCD-18Co PSAT1 OE cells, and EVs isolated from CCD-18Co PSAT1 OE cells. Quantification of TRAP-positive cell numbers in each group. The values are presented as the means ± SDs (n=5). N.S.: not significant by one-way ANOVA with Dunnett’s test.

**Figure S10. The effect of PSAT1 manipulation on EV characteristics. A.** Small RNA-seq analysis of MCF7 and BM02 (highly bone metastatic derivative of MCF7) breast cancer cells under PSAT1 manipulations. The differentially-expressed miRNAs (fold change of at least 2, p < 0.05, and average count > 10) between PSAT1 OE and control MCF7 cells and between PSAT1 KD and control BM02 cells were selected. The Venn diagram on the left shows the overlap between upregulated miRNAs in MCF7 PSAT1 OE cells and downregulated miRNAs in BM02 PSAT1 KD cells. The Venn diagram on the right shows the overlap between downregulated miRNAs in MCF7 PSAT1 OE cells and upregulated miRNAs in BM02 PSAT1 KD cells. **B**. Western blot analysis of RANK and RANKL in cells (left panels) and of EV markers such as CD9, CD63, and FLOT-1 in EVs (right panels) in PSAT1-overexpressing cell lines. **C.** NTA was performed to compare particle sizes between PSAT1-overexpressing BM02 cells and control BM02 cells. N.S.: not significant by Student’s t test.

**Figure S11. The effect of PSAT1 overexpression on bone metastasis in a breast cancer model. A.** IVIS imaging of mice injected with PSAT1-overexpressing BM02 and control BM02 cells on days 0, 7, and 28. **B.** Photon counting of BM02 cells, which express luciferase construct, in mice injected with PSAT1-overexpressing BM02 and control BM02 cells. The values are presented as the means ± SDs (n=5). *p<0.05 by Student’s t test. **C, D.** Images of HE staining (**C**) and TRAP staining (**D**) in bone from mice injected with PSAT1-overexpressing BM02 and control BM02 cells. Mice were sacrificed on day 28 after IVIS imaging. Top panels: low magnification and bottom panels: high magnification. Scale bars: 1 mm for the top panels and 100 μm for the bottom panels. **E**. Computed tomography of bones in mice injected with PSAT1-overexpressing BM02 and control BM02 cells. **F**. The effect of pretreatment with EVs to MCF7 cells on bone metastasis. MCF7 cell-derived EVs or MCF7 PSAT1 OE cell-derived EVs were isolated from the same amount of conditioned medium (11 mL) by a conventional ultracentrifugation method. MCF7 cells were pretreated with MCF7 cell-derived EVs or MCF7 PSAT1 OE cell-derived EVs for 2 weeks. As a negative control, MCF7 cells were treated with PBS. A total of 5 x 10^5^ MCF7 cells were injected via the caudal artery to establish the bone metastasis model. The metastatic cancer cells were monitored weekly by IVIS imaging.

