## supplemental figures for "Aberrant regulation of serine metabolism drives extracellular vesicle release and cancer progression"

### Fig. S1

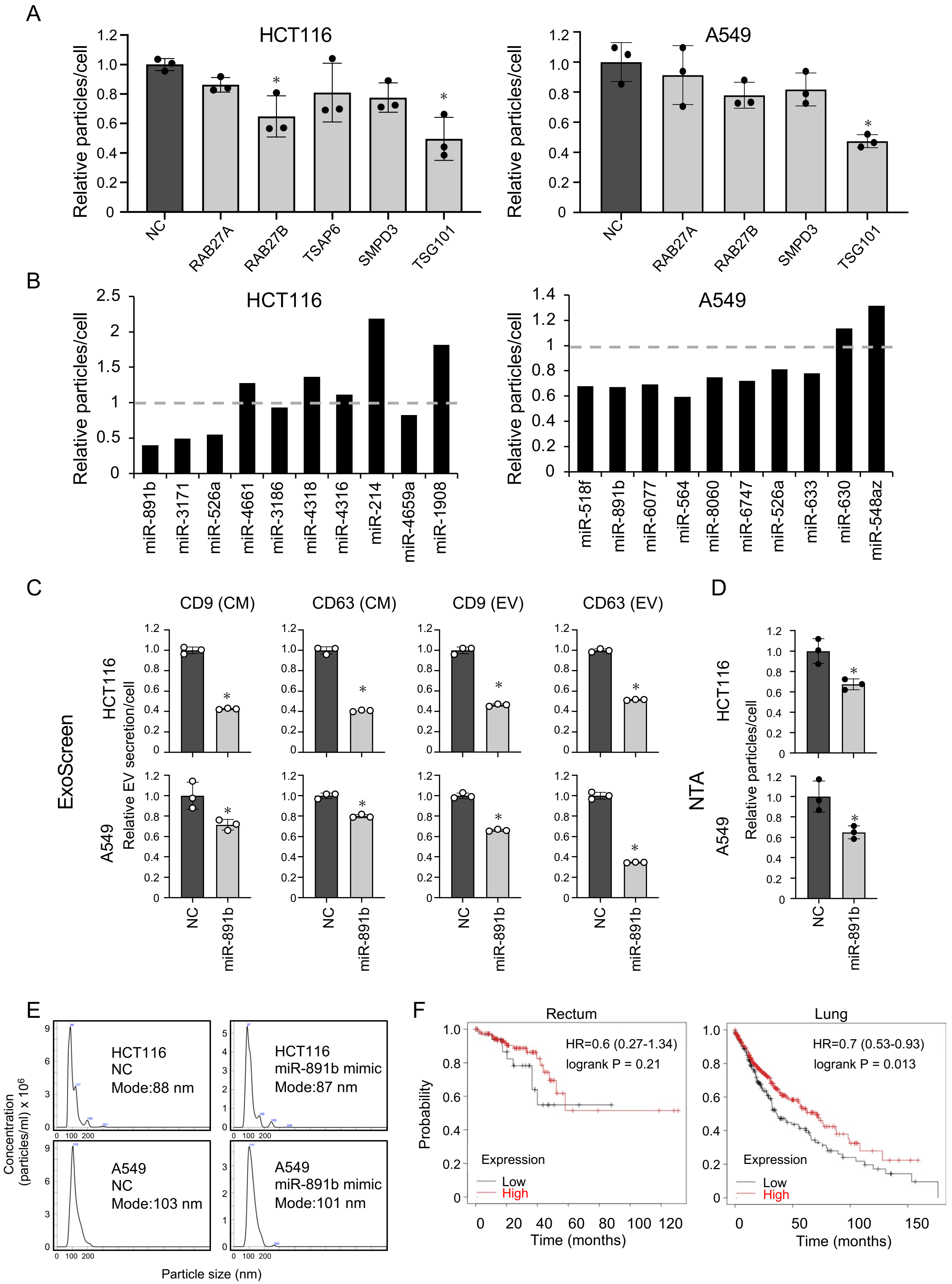

A
HCT116
A549
Relative particles/cell
NC
RAB27A
RAB27B
TSAP6
SMPD3
TSG101
B
HCT116
A549
miR-891b
miR-3171
miR-526a
miR-4661
miR-3186
miR-4318
miR-4316
miR-214
miR-4659a
miR-1908
miR-518f
miR-6077
miR-564
miR-8060
miR-6747
miR-633
miR-630
miR-548az
C
CD9 (CM)
CD63 (CM)
CD9 (EV)
CD63 (EV)
ExoScreen
Relative EV secretion/cell
D
NTA
E
HCT116
NC
Mode:88 nm
miR-891b mimic
Mode:87 nm
A549
Mode:103 nm
Mode:101 nm
Concentration (particles/ml) x 10^6
Particle size (nm)
F
Rectum
Lung
HR=0.6 (0.27-1.34)
logrank P = 0.21
HR=0.7 (0.53-0.93)
logrank P = 0.013
Probability
Expression
Low
High
Time (months)

### Fig. S2

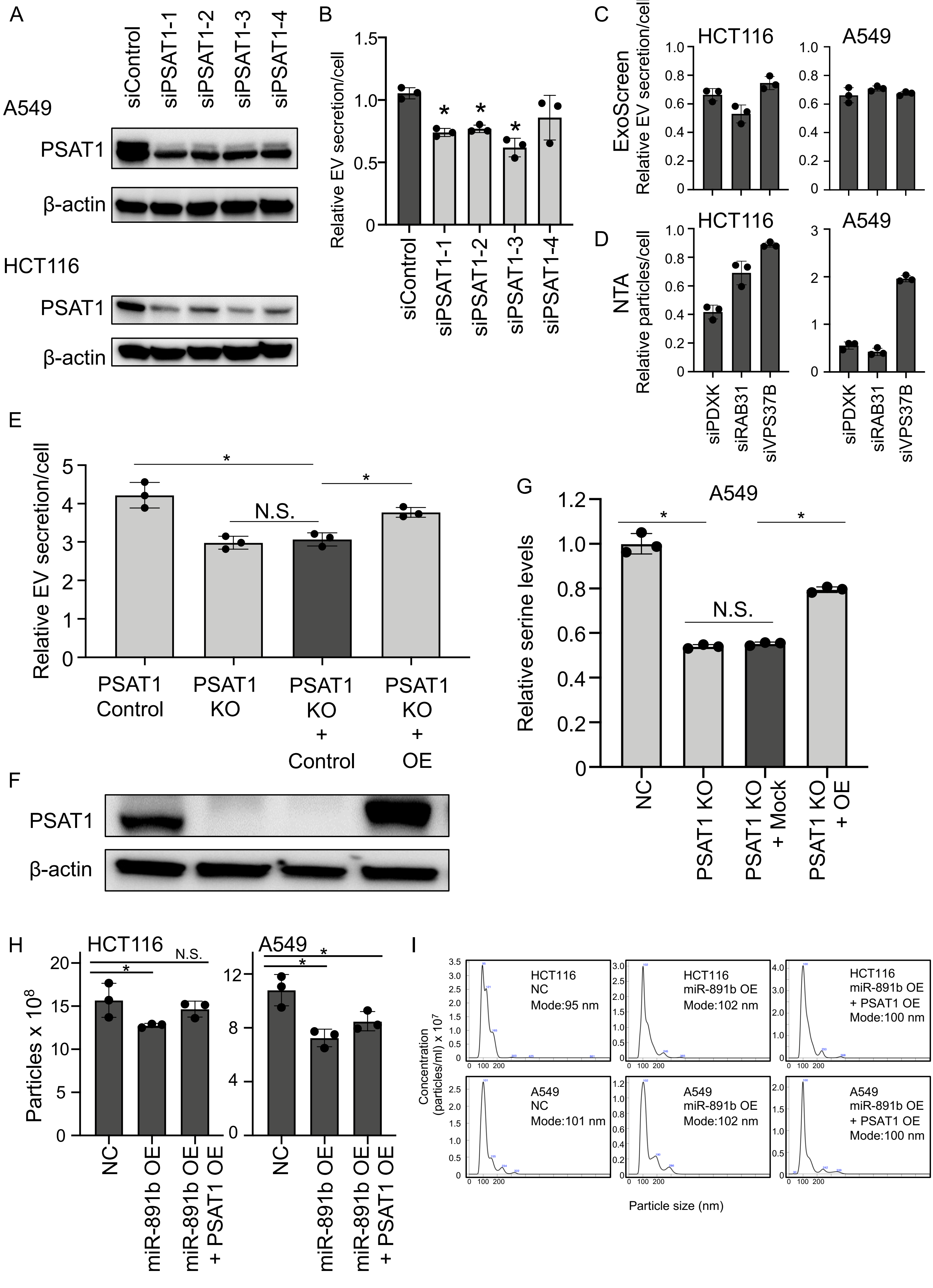
A
siControl
siPSAT1-1
siPSAT1-2
siPSAT1-3
siPSAT1-4
A549
PSAT1
β-actin
HCT116
PSAT1
β-actin
B
Relative EV secretion/cell
siControl
siPSAT1-1
siPSAT1-2
siPSAT1-3
siPSAT1-4
C
ExoScreen
Relative EV secretion/cell
HCT116
A549
D
NTA
Relative particles/cell
HCT116
A549
siPDXK
siRAB31
siVPS37B
E
Relative EV secretion/cell
N.S.
PSAT1 Control
PSAT1 KO
PSAT1 KO + Control
PSAT1 KO + OE
F
PSAT1
β-actin
G
A549
Relative serine levels
N.S.
NC
PSAT1 KO
PSAT1 KO + Mock
PSAT1 KO + OE
H
HCT116
A549
N.S.
Particles x 10^8
NC
miR-891b OE
miR-891b OE + PSAT1 OE
I
HCT116 NC Mode:95 nm
HCT116 miR-891b OE Mode:102 nm
HCT116 miR-891b OE + PSAT1 OE Mode:100 nm
A549 NC Mode:101 nm
A549 miR-891b OE Mode:102 nm
A549 miR-891b OE + PSAT1 OE Mode:100 nm
Concentration (particles/ml) x 10^7
Particle size (nm)

### Fig. S3

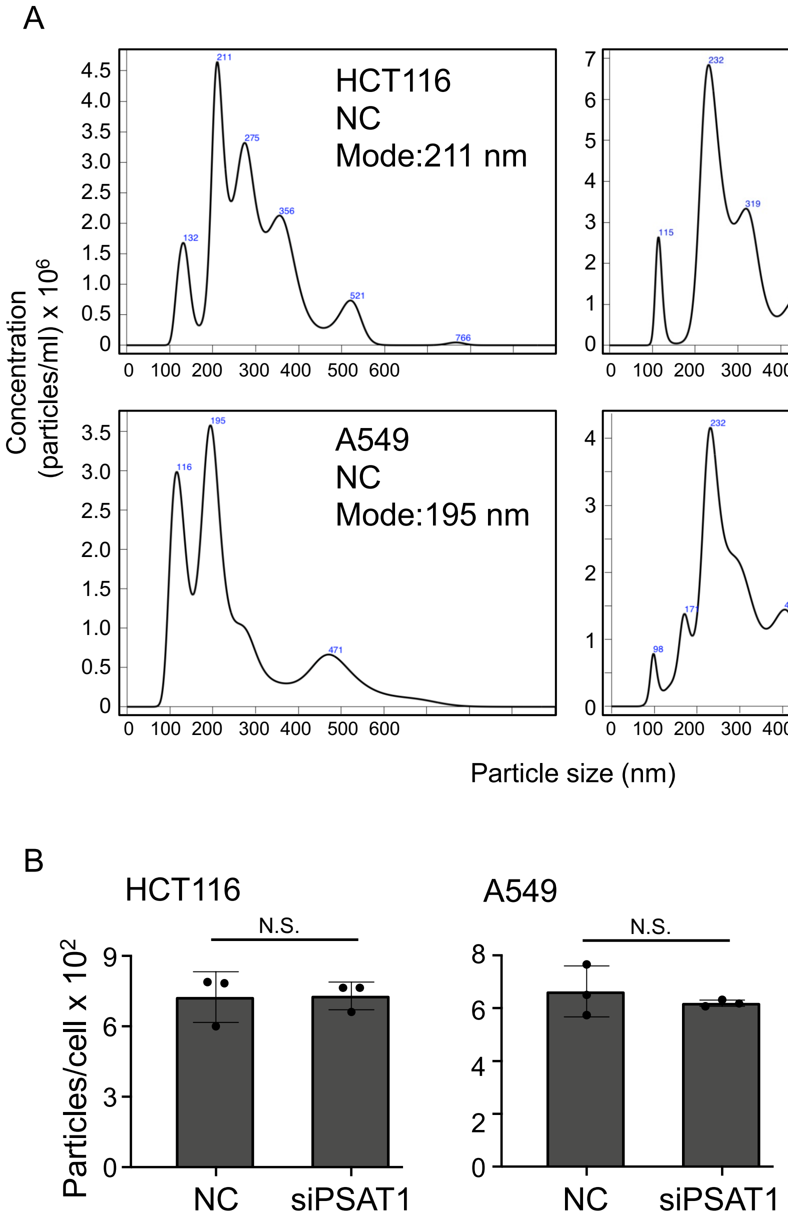
A
HCT116
NC
Mode:211 nm
HCT116
siPSAT1
Mode:232 nm
A549
NC
Mode:195 nm
A549
siPSAT1
Mode:232 nm
Concentration
(particles/ml) x 10^6
Particle size (nm)
B
HCT116
A549
N.S.
Particles/cell x 10^2
NC
siPSAT1

Fig. S4

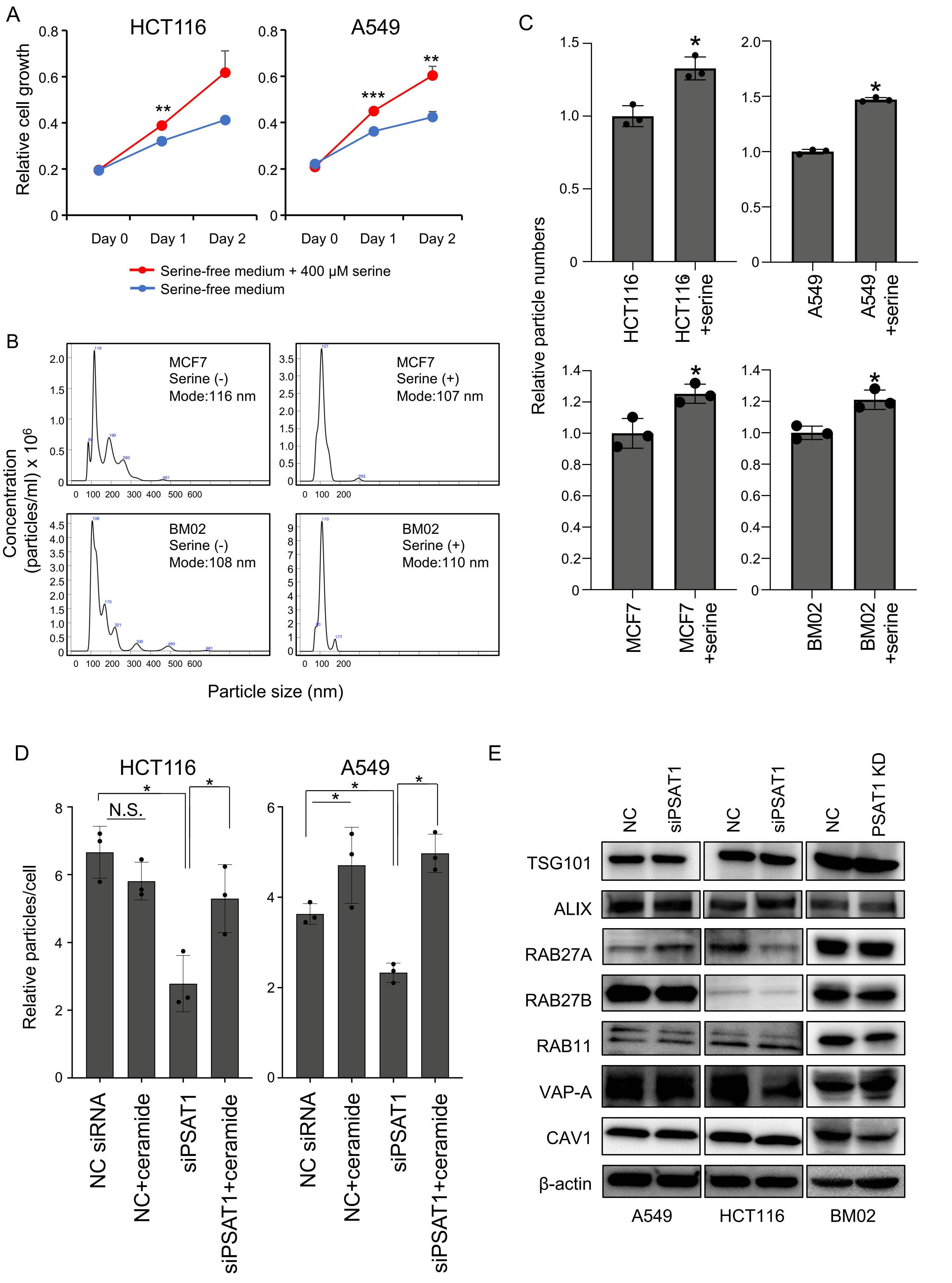

A
HCT116
A549
Relative cell growth
Day 0
Day 1
Day 2
Serine-free medium + 400 μM serine
Serine-free medium
B
MCF7
Serine (-)
Mode:116 nm
MCF7
Serine (+)
Mode:107 nm
BM02
Serine (-)
Mode:108 nm
BM02
Serine (+)
Mode:110 nm
Concentration (particles/ml) x 10^6
Particle size (nm)
C
Relative particle numbers
HCT116
HCT116 +serine
A549
A549 +serine
MCF7
MCF7 +serine
BM02
BM02 +serine
D
HCT116
A549
N.S.
Relative particles/cell
NC siRNA
NC+ceramide
siPSAT1
siPSAT1+ceramide
E
NC
siPSAT1
NC
siPSAT1
NC
PSAT1 KD
TSG101
ALIX
RAB27A
RAB27B
RAB11
VAP-A
CAV1
β-actin
A549
HCT116
BM02

### Fig. S5

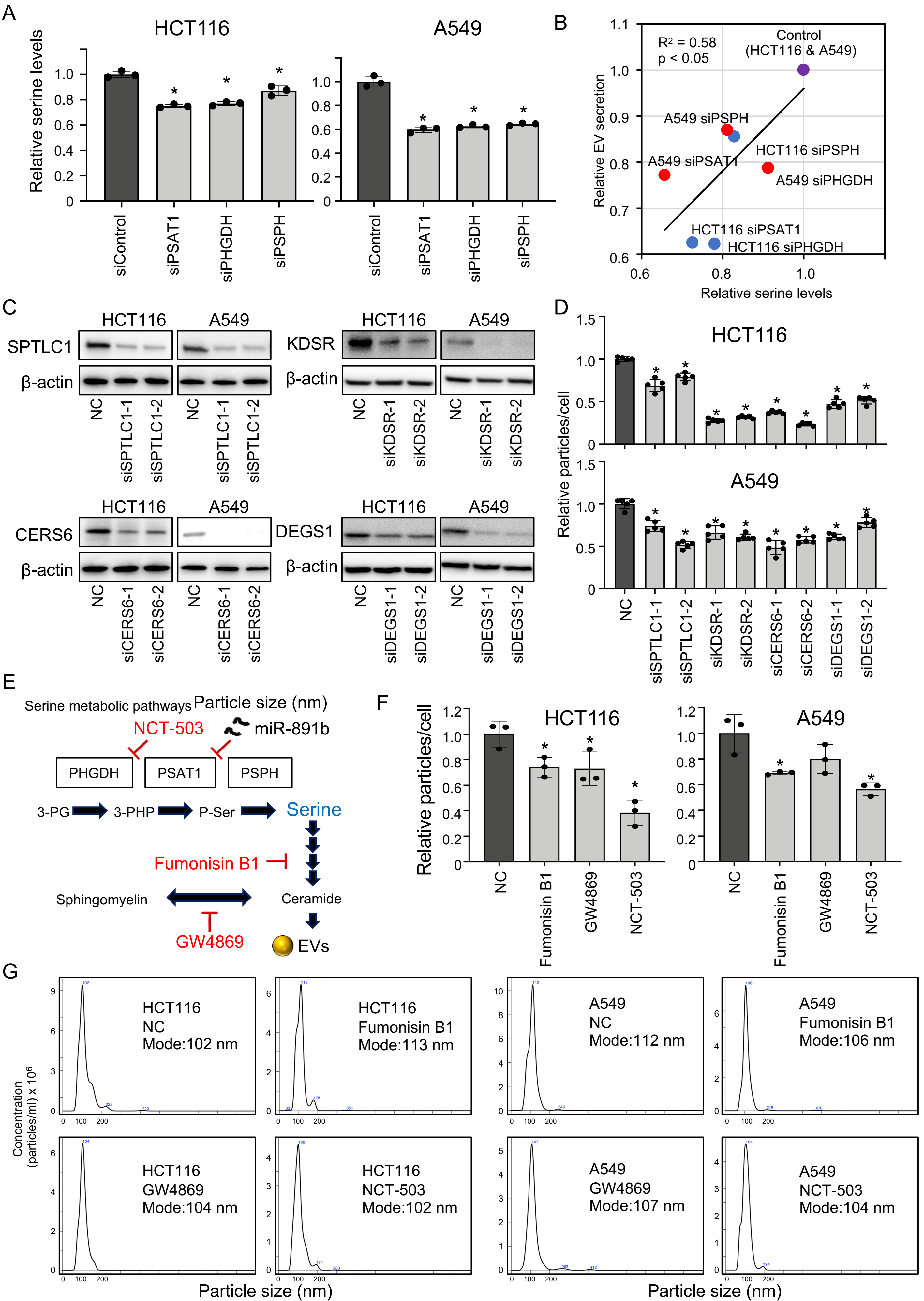
A
HCT116
A549
Relative serine levels
siControl
siPSAT1
siPHGDH
siPSPH
B
R² = 0.58
p < 0.05
Control (HCT116 & A549)
A549 siPSPH
A549 siPSAT1
HCT116 siPSPH
A549 siPHGDH
HCT116 siPSAT1
HCT116 siPHGDH
Relative EV secretion
Relative serine levels
C
SPTLC1
KDSR
CERS6
DEGS1
β-actin
NC
siSPTLC1-1
siSPTLC1-2
siKDSR-1
siKDSR-2
siCERS6-1
siCERS6-2
siDEGS1-1
siDEGS1-2
D
Relative particles/cell
E
Serine metabolic pathways
NCT-503
miR-891b
PHGDH
PSAT1
PSPH
3-PG
3-PHP
P-Ser
Serine
Fumonisin B1
Sphingomyelin
Ceramide
GW4869
EVs
F
Relative particles/cell
Fumonisin B1
GW4869
NCT-503
G
Concentration (particles/ml) x 10⁶
HCT116 NC Mode:102 nm
HCT116 Fumonisin B1 Mode:113 nm
A549 NC Mode:112 nm
A549 Fumonisin B1 Mode:106 nm
HCT116 GW4869 Mode:104 nm
HCT116 NCT-503 Mode:102 nm
A549 GW4869 Mode:107 nm
A549 NCT-503 Mode:104 nm
Particle size (nm)

Fig. S6

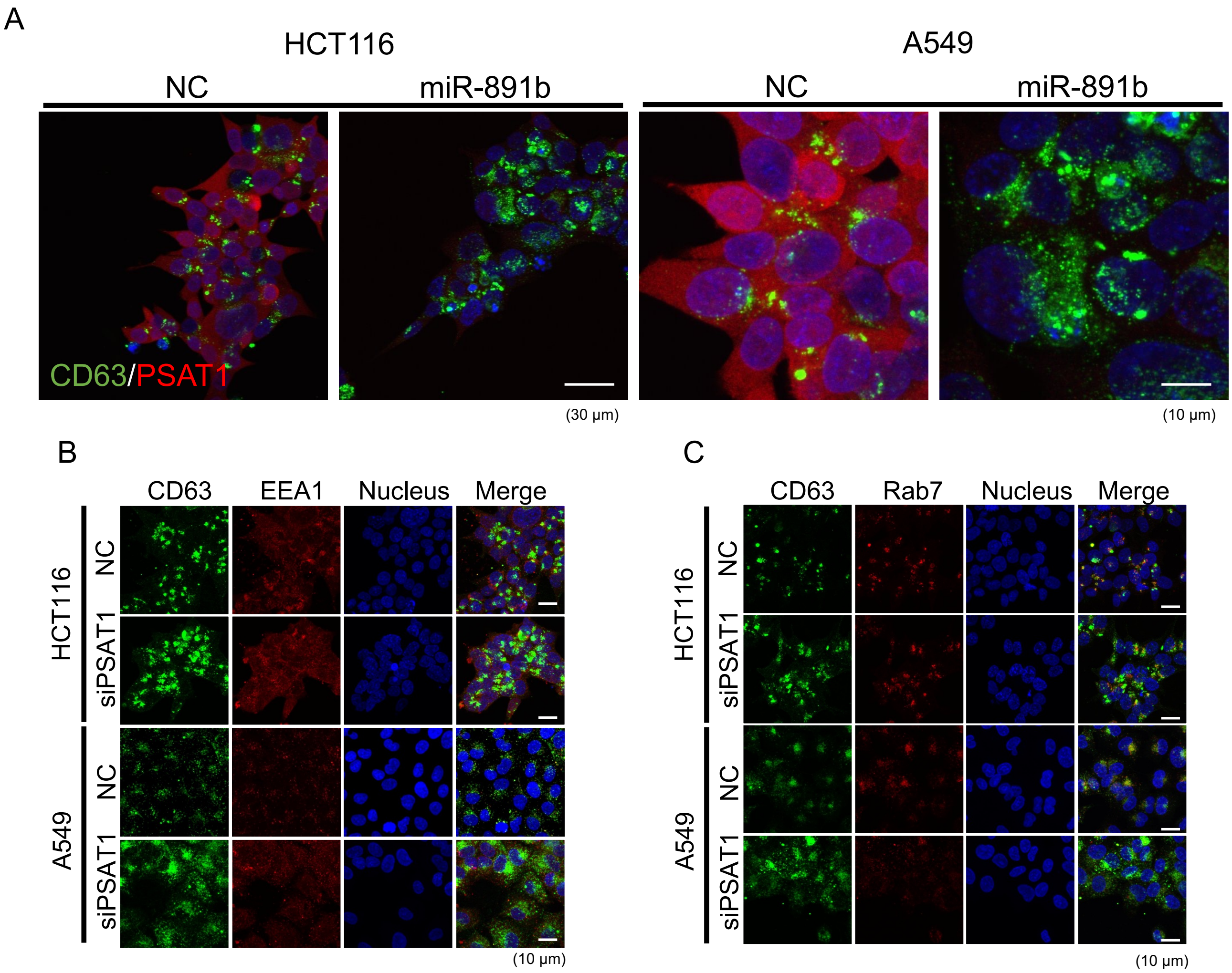

A
HCT116
A549
NC
miR-891b
NC
miR-891b
CD63/PSAT1
(30 μm)
(10 μm)
B
CD63
EEA1
Nucleus
Merge
HCT116
NC
siPSAT1
A549
NC
siPSAT1
(10 μm)
C
CD63
Rab7
Nucleus
Merge
HCT116
NC
siPSAT1
A549
NC
siPSAT1
(10 μm)

### Fig. S7

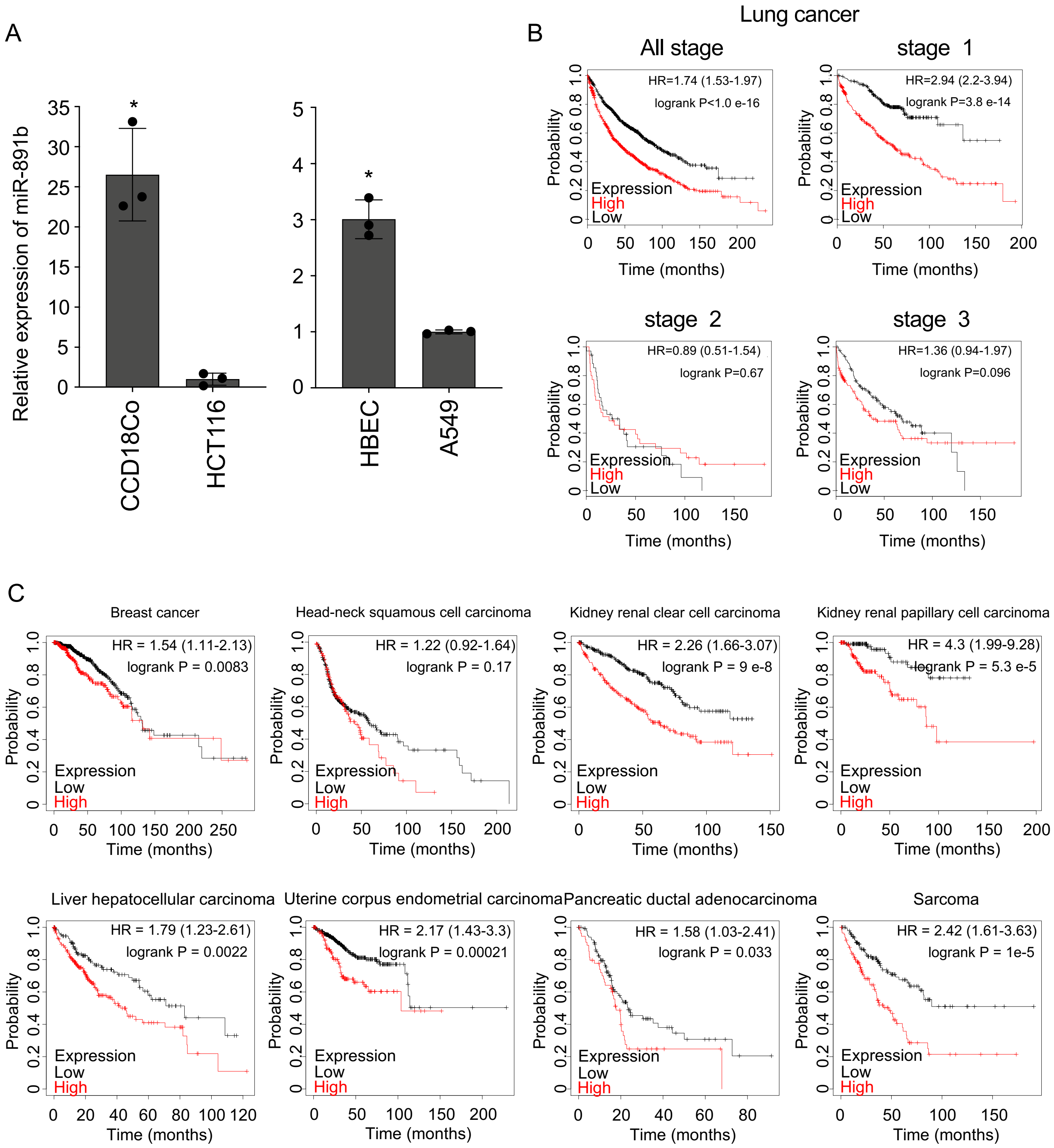

A
Relative expression of miR-891b
CCD18Co
HCT116
HBEC
A549
B
Lung cancer
All stage
HR=1.74 (1.53-1.97)
logrank P<1.0 e-16
stage 1
HR=2.94 (2.2-3.94)
logrank P=3.8 e-14
stage 2
HR=0.89 (0.51-1.54)
logrank P=0.67
stage 3
HR=1.36 (0.94-1.97)
logrank P=0.096
Expression
High
Low
Probability
Time (months)
C
Breast cancer
HR = 1.54 (1.11-2.13)
logrank P = 0.0083
Head-neck squamous cell carcinoma
HR = 1.22 (0.92-1.64)
logrank P = 0.17
Kidney renal clear cell carcinoma
HR = 2.26 (1.66-3.07)
logrank P = 9 e-8
Kidney renal papillary cell carcinoma
HR = 4.3 (1.99-9.28)
logrank P = 5.3 e-5
Liver hepatocellular carcinoma
HR = 1.79 (1.23-2.61)
logrank P = 0.0022
Uterine corpus endometrial carcinoma
HR = 2.17 (1.43-3.3)
logrank P = 0.00021
Pancreatic ductal adenocarcinoma
HR = 1.58 (1.03-2.41)
logrank P = 0.033
Sarcoma
HR = 2.42 (1.61-3.63)
logrank P = 1e-5
Expression
Low
High
Probability
Time (months)

Fig. S8

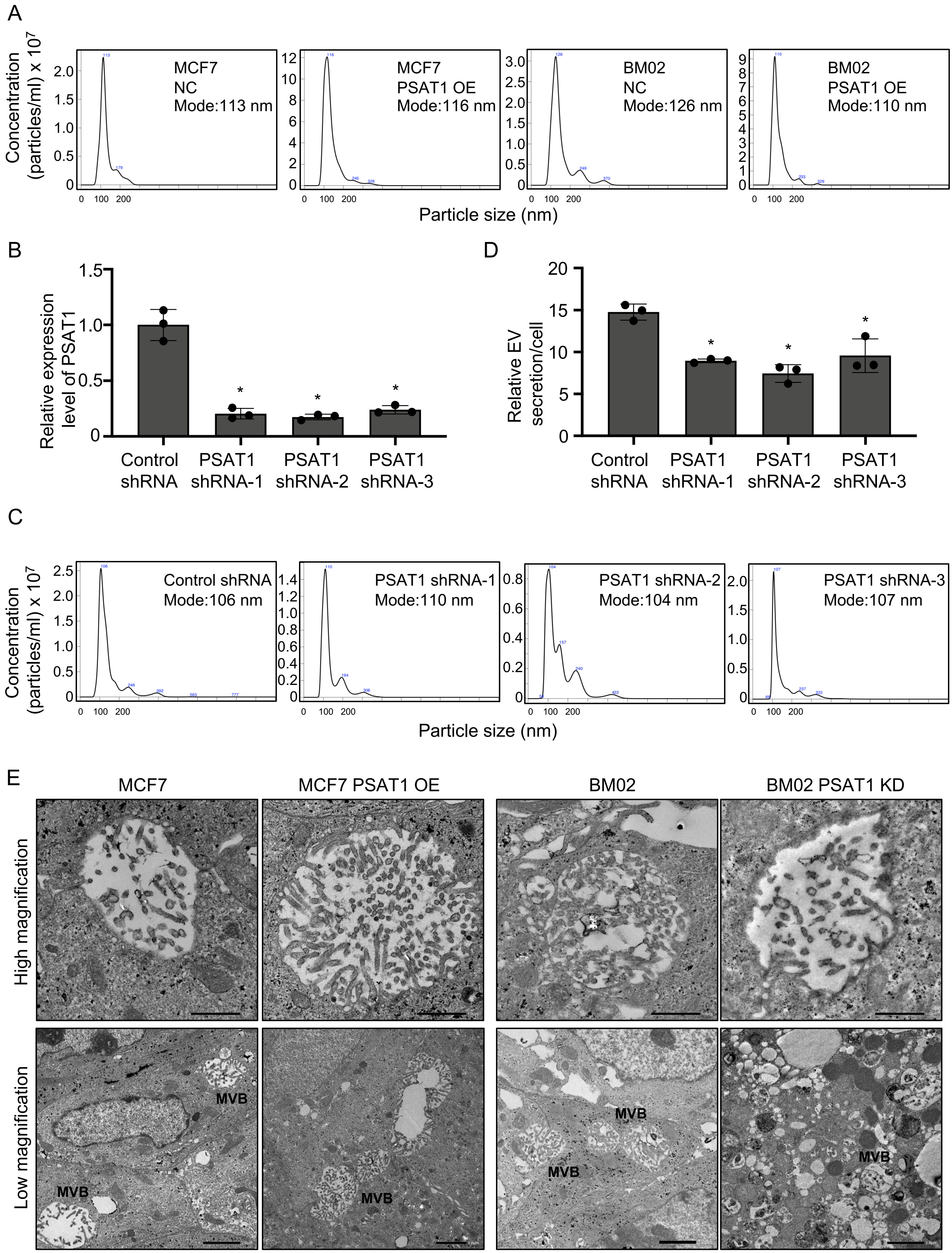
A
Concentration (particles/ml) x 10^7
MCF7
NC
Mode:113 nm
MCF7
PSAT1 OE
Mode:116 nm
BM02
NC
Mode:126 nm
BM02
PSAT1 OE
Mode:110 nm
Particle size (nm)
B
Relative expression level of PSAT1
Control shRNA
PSAT1 shRNA-1
PSAT1 shRNA-2
PSAT1 shRNA-3
D
Relative EV secretion/cell
Control shRNA
PSAT1 shRNA-1
PSAT1 shRNA-2
PSAT1 shRNA-3
C
Concentration (particles/ml) x 10^7
Control shRNA
Mode:106 nm
PSAT1 shRNA-1
Mode:110 nm
PSAT1 shRNA-2
Mode:104 nm
PSAT1 shRNA-3
Mode:107 nm
Particle size (nm)
E
MCF7
MCF7 PSAT1 OE
BM02
BM02 PSAT1 KD
High magnification
Low magnification
MVB

Fig. S9

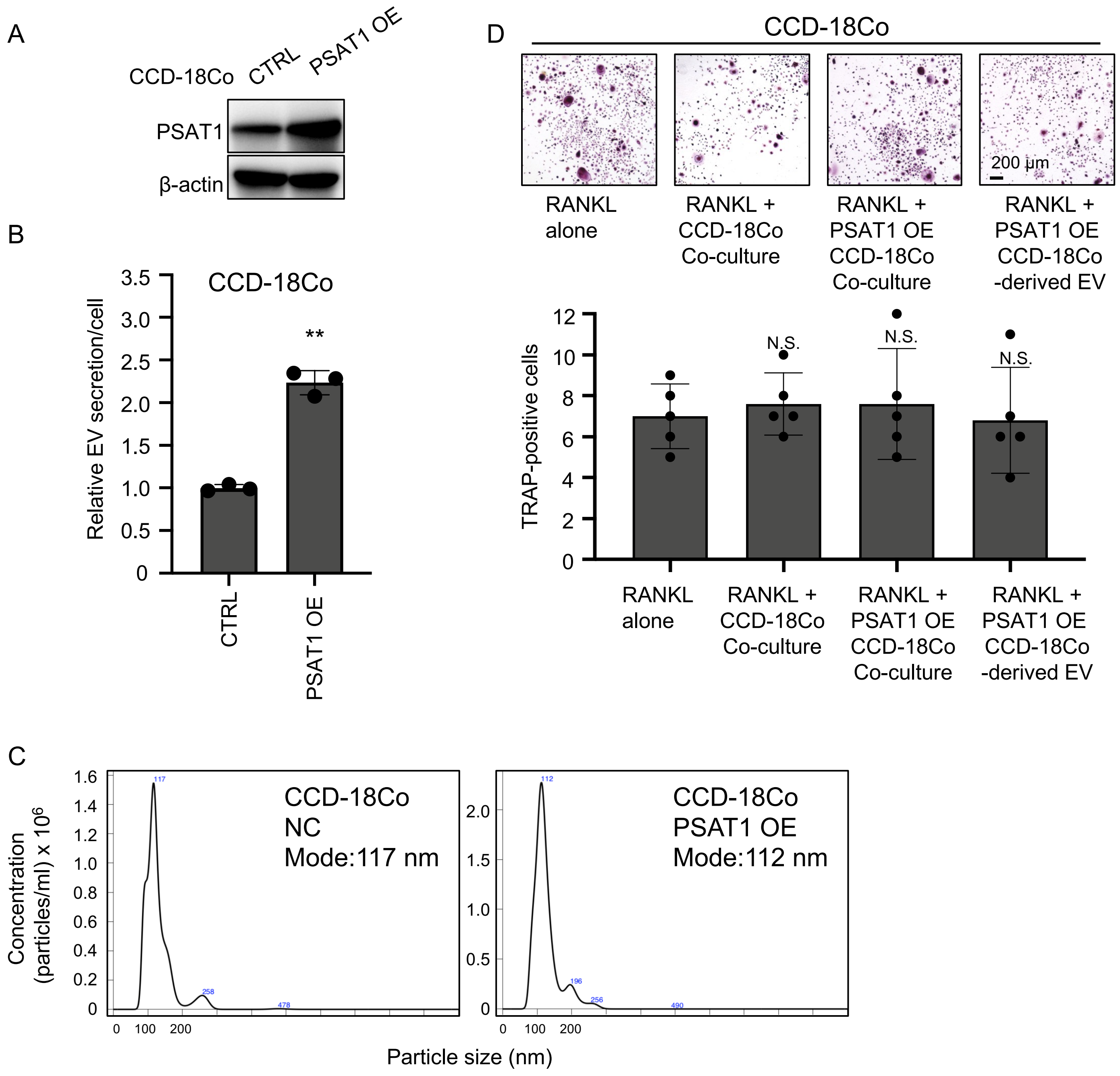

A
CCD-18Co
CTRL
PSAT1 OE
PSAT1
β-actin
B
CCD-18Co
Relative EV secretion/cell
3.5
3.0
2.5
2.0
1.5
1.0
0.5
0
**
CTRL
PSAT1 OE
C
Concentration (particles/ml) x 10^6
CCD-18Co
NC
Mode:117 nm
CCD-18Co
PSAT1 OE
Mode:112 nm
Particle size (nm)
0
100
200
D
CCD-18Co
200 µm
RANKL alone
RANKL + CCD-18Co Co-culture
RANKL + PSAT1 OE CCD-18Co Co-culture
RANKL + PSAT1 OE CCD-18Co -derived EV
TRAP-positive cells
12
10
8
6
4
2
0
N.S.
N.S
N.S.

### Fig. S10

A

MCF7 PSAT1 OE

OE vs. Control
2 fold-or-more
$p < 0.05$
average count > 10

11 miRNAs: high
1 miRNA: low

BM02 PSAT1 KD

KD vs. Control
2 fold-or-more
$p < 0.05$
average count > 10

31 miRNAs: high
0 miRNA: low

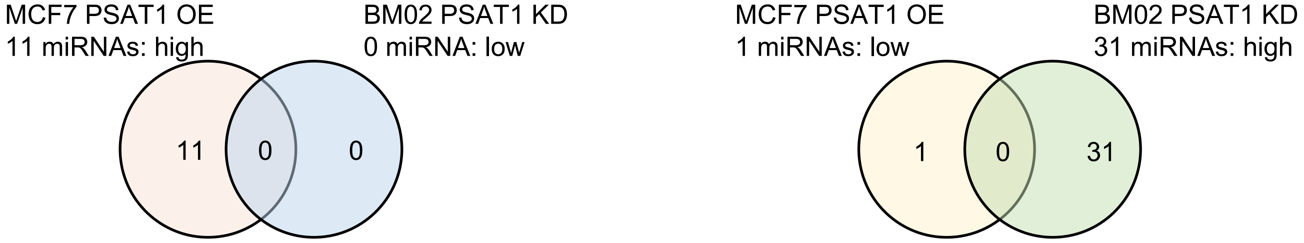


B

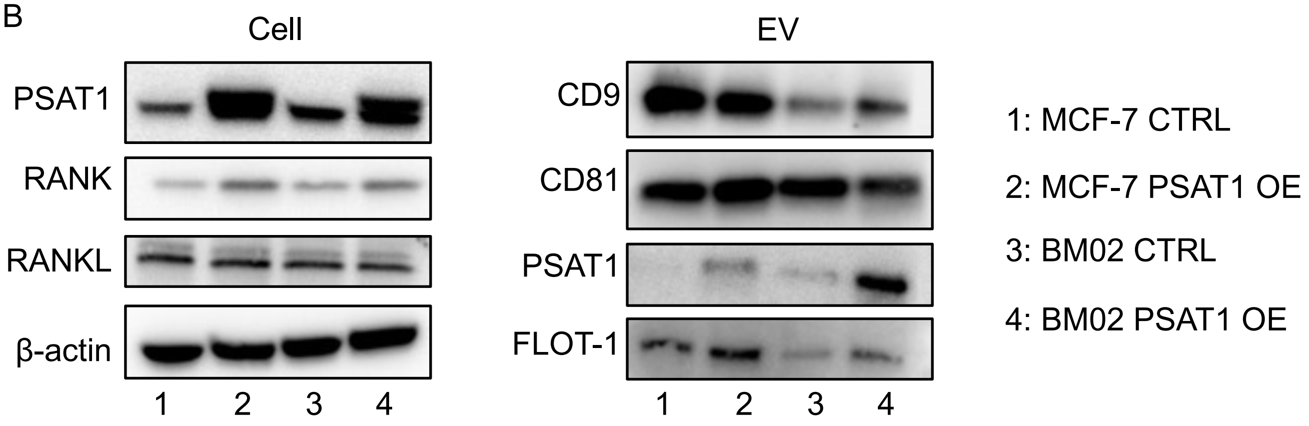


C

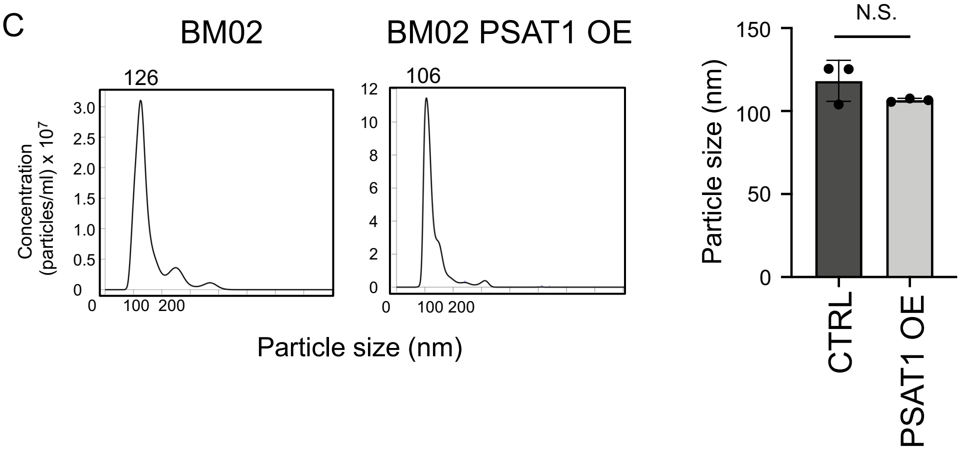

### Fig. S11

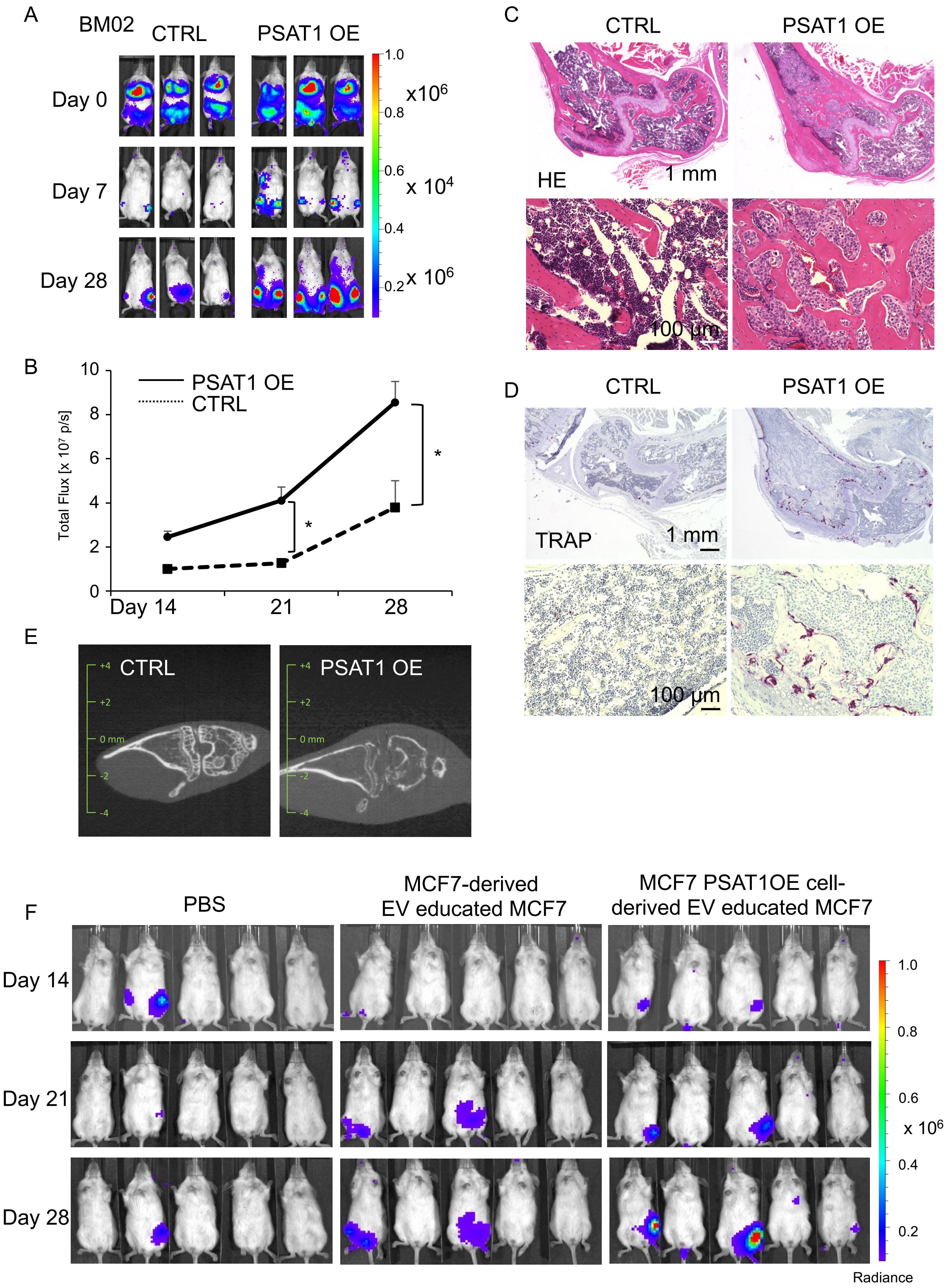

A
BM02
CTRL
PSAT1 OE
Day 0
Day 7
Day 28
1.0
0.8
0.6
0.4
0.2
x10^6
x 10^4
x 10^6
B
PSAT1 OE
CTRL
Total Flux [x 10^7 p/s]
10
8
6
4
2
0
Day 14
21
28
*
*
C
CTRL
PSAT1 OE
HE
1 mm
100 µm
D
CTRL
PSAT1 OE
TRAP
1 mm
100 µm
E
CTRL
PSAT1 OE
+4
+2
0 mm
-2
-4
F
PBS
MCF7-derived EV educated MCF7
MCF7 PSAT1OE cell-derived EV educated MCF7
Day 14
Day 21
Day 28
1.0
0.8
0.6
0.4
0.2
x 10^6
Radiance
